# NISNet3D: Three-Dimensional Nuclear Synthesis and Instance Segmentation for Fluorescence Microscopy Images

**DOI:** 10.1101/2022.06.10.495713

**Authors:** Liming Wu, Alain Chen, Paul Salama, Kenneth Dunn, Edward Delp

## Abstract

The primary step in tissue cytometry is the automated distinction of individual cells (segmentation). Since cell borders are seldom labeled, researchers generally segment cells by their nuclei. While effective tools have been developed for segmenting nuclei in two dimensions, segmentation of nuclei in three-dimensional volumes remains a challenging task for which few tools have been developed. The lack of effective methods for three-dimensional segmentation represents a bottleneck in the realization of the potential of tissue cytometry, particularly as methods of tissue clearing present researchers with the opportunity to characterize entire organs. Methods based upon deep-learning have shown enormous promise, but their implementation is hampered by the need for large amounts of manually annotated training data. In this paper we describe 3D Nuclei Instance Segmentation Network (NISNet3D), a deep learning-based approach in which training is accomplished using synthetic data, profoundly reducing the effort required for network training. We compare results obtained from NISNet3D with results obtained from eight existing techniques.

## 1 Introduction

Over the past ten years, various technological developments have provided biologists with the ability to collect microscopy images of enormous scale and complexity. Methods of tissue clearing combined with automated confocal or lightsheet microscopes have enabled three-dimensional imaging of entire organs or even entire organisms at subcellular resolution. Novel methods of multiplexing have been developed so that researchers can now simultaneously characterize 50 or more targets in the same tissue. However, as biologists turn to the task of analyzing these extraordinary volumes (tissue cytometry), they quickly discover that the methods of automated image analysis necessary for extracting quantitative data from images of this scale are frequently inadequate for the task. In particular, while effective methods for distinguishing (segmenting) individual cells are available for analyses of two-dimensional images, corresponding methods for segmenting cells in three-dimensional volumes are generally lacking. The problem of three-dimensional image segmentation thus represents a bottleneck in the full realization of 3D tissue cytometry as a tool in biological microscopy.

There are two main categories of segmentation approaches, non-machinelearning image processing and computer vision techniques and techniques based on machine learning and in particular deep learning [1, 2]. The traditional image processing techniques (e.g. watershed, thresholding, edge detection, and morphological operations) can be effective on one type of images but may not generalize to other types of images without careful parameter tuning. Segmentation techniques based on deep learning have shown great promise, in some cases providing accurate and robust results across a range of image types [3–7]. However, their utility is limited by the large amounts of manually annotated (ground truth) data needed for training, validation, and testing. Annotation is a labor-intensive and time-consuming process, especially for a 3D volume. While tools have been developed to facilitate the laborious process of manual annotation [8–11], the generation of training data for microscopy images remains a major obstacle to implementing segmentation approaches based upon deep learning.

The problem of generating sufficient training data can be alleviated using data augmentation, a process in which existing manually annotated training data is supplemented with synthetic data generated from modifications of the manually annotated data [12–15]. An alternative approach is to use synthetic data for training [14, 16, 17]. A method described in [18] generates synthetic 3D microscopy volumes by stacking 2D synthetic image slices using 2D distributions of fluorescent markers that are generated using Generative Adversarial Networks (GANs). Similarly, a 3D GAN was used in [19] to generate fully 3D volumetric cell masks. We have shown [7, 20–22] that GANs can be used to generate synthetic microscopy volumes that can be used for training, and incorporated this approach into the DeepSynth segmentation system [6].

Convolutional Neural Networks (CNNs) have had great success for solving problems such as object classification, detection, and segmentation [23, 24]. The encoder-decoder architecture has been widely used for biomedical image analysis including volumetric segmentation [25–27], medical image registration [28], and nuclear segmentation [3, 6, 15, 29–33]. However, most CNNs are designed for segmenting two-dimensional images and cannot be directly used for segmentation of 3D volumes [29, 32, 34]. Other methods described in [3, 14, 30] process images slice by slice and fuse together two dimensional results to form 3D segmentation results that fail to represent the 3D anisotropy of microscopy volumes.

In this paper we describe the 3D Nuclei Instance Segmentation Network (NISNet3D), a deep learning-based segmentation technique that can use synthetic volumes, manually annotated volumes, or a combination of synthetic and annotated volumes. NISNet3D is a true three-dimensional segmentation method that operates directly on 3D volumes, using 3D CNNs to exploit 3D information in a microscopy volume, thereby generating more accurate segmentations of nuclei in 3D image volumes.

## 2 Results

### 2.1 3D Nuclei Image Synthesis

Deep learning methods generally require large amounts of training samples to achieve accurate results. However, manually annotating ground truth is a tedious task and impractical in many situations especially for 3D microscopy volumes. To address this issue, we generate synthetic microscopy volumes for training NISNet3D. In particular, we enhance the synthetic microscopy volume generation method described in [6, 7, 31] to model non-ellipsoidal or irregularly shaped nuclei, in contrast to the prior methods described in [6, 7] which model nuclei as regular ellipsoids using the approach described in [31]. It must be emphasized that NISNet3D can use synthetic volumes or annotated real volumes or combinations of both. We will demonstrate this in our experiments. For generating microscopy volumes, we first generate synthetic segmentation masks, which will be used as the ground truth masks during training, by iteratively adding initial binary nuclei to an empty 3D volume. The details for generating initial binary nuclei are described in Section 3.2.1. Since many of the nuclei in the original microscopy images are not strictly ellipsoidal and look more like deformed ellipsoids (See Figure 2 (left)), We model these initial nuclei as deformed 3D binary ellipsoids with random size and orientation. Specifically, we use elastic transformation [35] to deform the 3D binary mask of the nuclei. The details for deforming the initial binary nuclei are described in Section 3.2.1.

Examples of deformed ellipsoids are shown in Figure 2 (right column). We then use the unpaired image-to-image translation model known as SpCycleGAN [7, 36] for generating synthetic microscopy volumes. By unpaired we mean that the binary segmentation mask we created above is not the ground truth of actual microscopy images. The input to the SpCycleGAN is the binary segmentation masks we created and actual microscopy images (i.e. unpaired). As shown in Figure 1, we use the binary segmentation mask we created and actual microscopy volumes for training the SpCycleGAN. After training we generate synthetic microscopy volumes by using different synthetic microscopy segmentation masks we created as input to the SpCycleGAN. Note that since the SpCycleGAN generates 2D slices, we sequentially generate 2D slices for each focal plane and stack them to construct a 3D synthetic volume. During our experiments, we observed that if we only model nuclei as regular ellipsoids when the original microscopy volume contains many non-ellipsoidal or irregularly shaped nuclei, the SpCycleGAN will have difficulty learning the mapping function between the the binary masks and microscopy images, resulting in generating many non-ellipsoidal nuclei which are not aligned well with the corresponding binary masks. Including non-ellipsoidal nuclei or irregularly shaped nuclei will help SpCycleGAN learn a more accurate mapping and thus the SpCycleGAN can generate more representative synthetic images. We have generated 950 synthetic microscopy volumes in total, and each is of size 128×128×128 using the representative subvolumes of different datasets. The examples of actual microscopy images and corresponding synthetic microscopy images are shown in Figure 3.

**Fig. 1.**
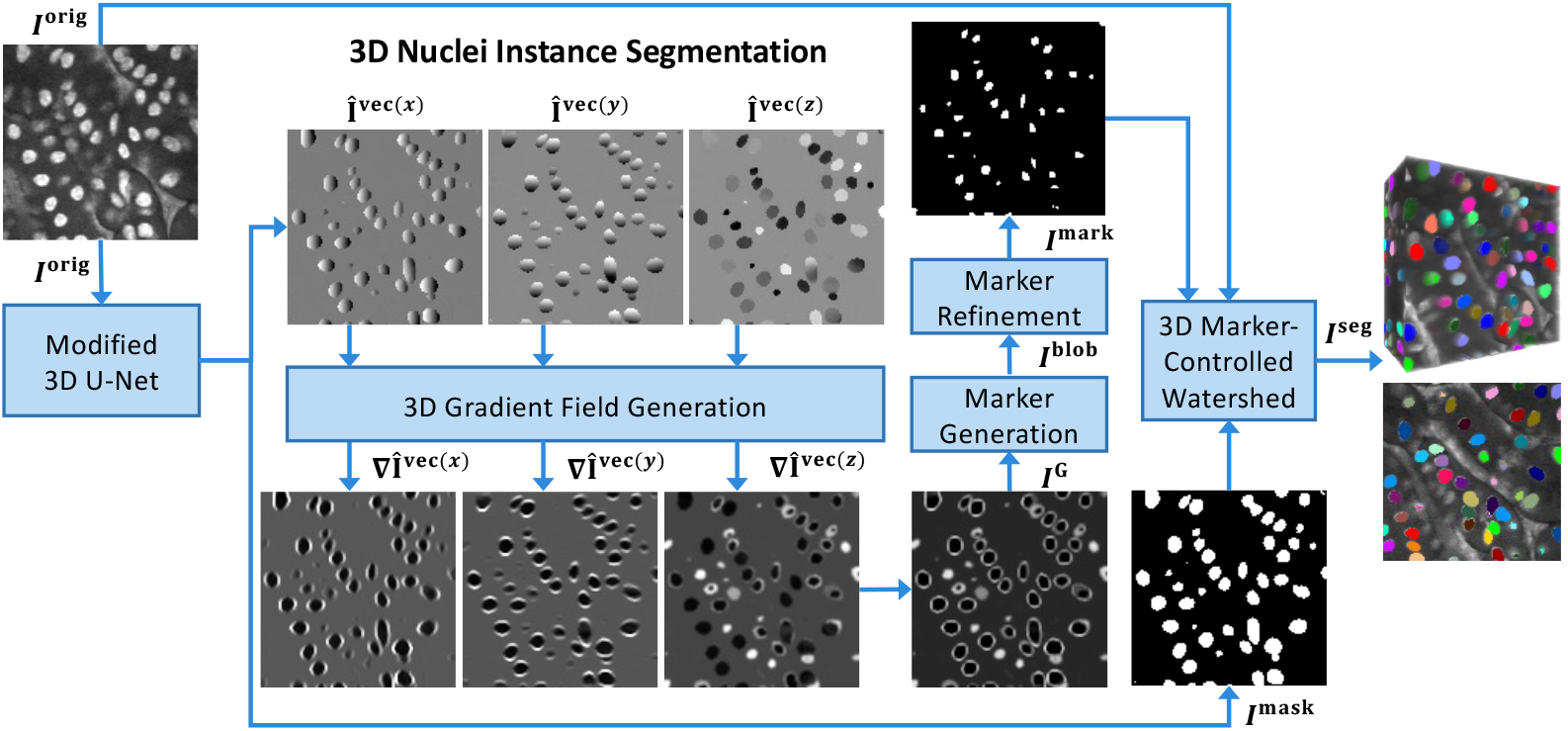
Overview of NISNet3D for 3D nuclei instance segmentation. NISNet3D uses a modified 3D U-Net for nuclei segmentation and 3D vector field array estimation where each voxel represents a 3D vector pointing to the nearest nuclei centroid. NISNet3D then generates a 3D gradient field array from the 3D vector field array and further generates refined markers for 3D watershed segmentation

**Fig. 2.**
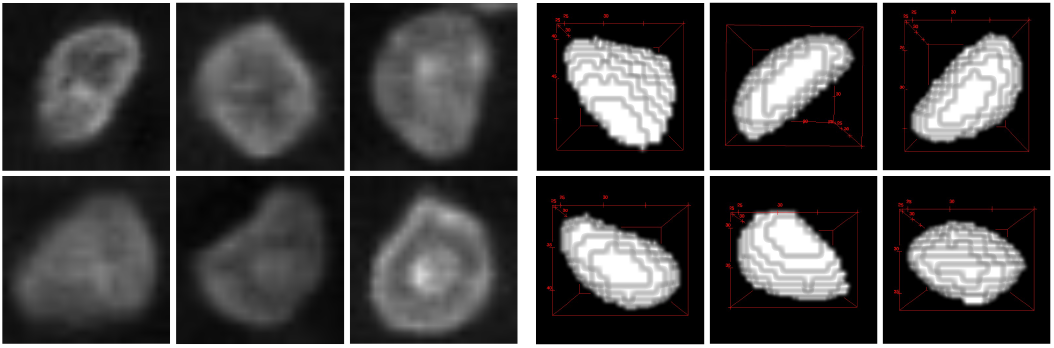
Non-ellipsoidal shaped nuclei in actual microscopy volumes (left column) and synthetic binary nuclei segmentation masks after using elastic deformation (right column)

**Fig. 3.**
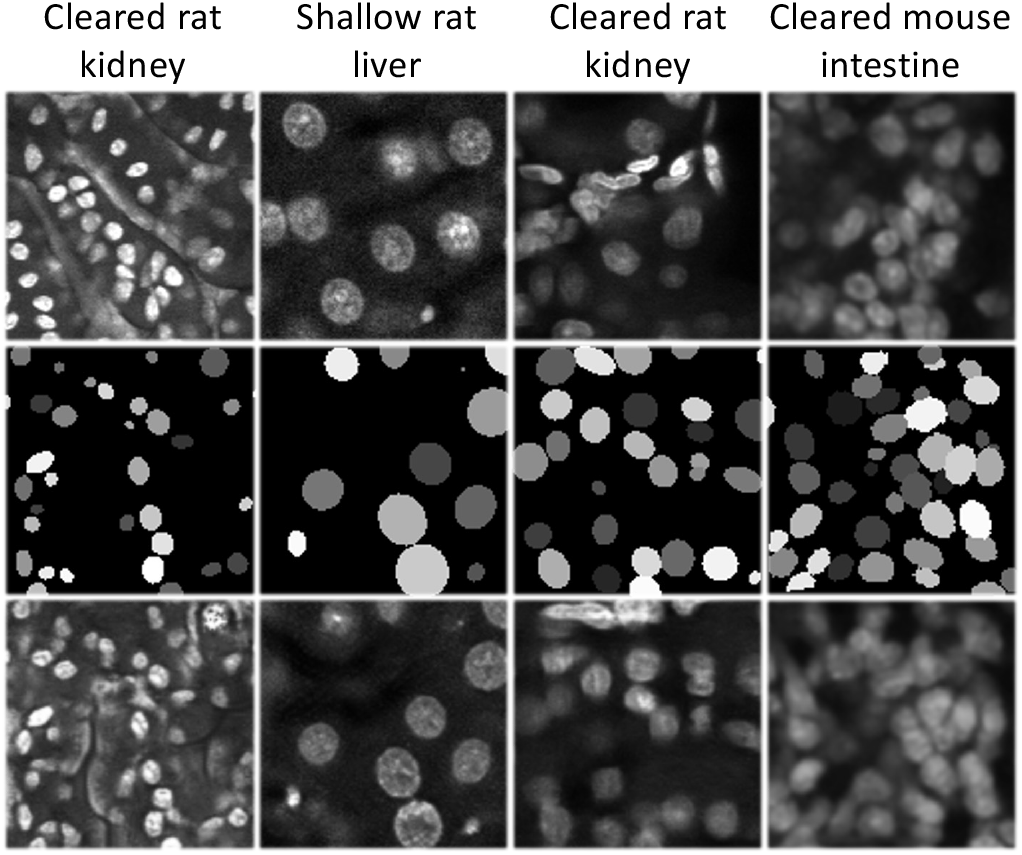
Actual microscopy images (top row), synthetic segmentation masks (middle row), and corresponding synthetic microscopy images (bottom row)

### 2.2 Evaluation Datasets

We used four microscopy volumes in our experiments, denoted by 𝒱_1_-𝒱_4_, having fluorescent-labeled (Hoechst 33342 stain) nuclei that were collected from cleared rat kidneys, shallow rat livers, and cleared mouse intestines using confocal microscopy, to both generate synthetic data as well as provide ground truth data. Datasets 𝒱_1_-𝒱_4_ are available at [37]. In particular, we use the 3D Nuclei Image Synthesis method described in Section 2.1 in conjunction with a subset of 𝒱_1_-𝒱_4_ (other than those used for validation and testing) to generate 950 volumes of corresponding synthetic microscopy volumes (250 volumes for 𝒱_1_, 250 volumes for 𝒱_2_, 250 volumes for 𝒱_3_, and 200 volumes for 𝒱_4_), that were then used for training NISNet3D. We also manually annotated representative subvolumes of 𝒱_1_-𝒱_4_ using ITK-SNAP [8], which are used as ground truth data. We have annotated 1, 16, 9, 5 subvolumes for 𝒱_1_-𝒱_4_, respectively. It is to be noted that both 𝒱_1_ and 𝒱_3_ are cleared rat kidney volumes but show different nuclear morphology. In addition, we also used a publicly available electron microscopy zebrafish brain volume, 𝒱_5_, known as NucMM [38], to evaluate NISNet3D. We use the NucMM dataset to demonstrate that NISNet3D not only works on fluorescent microscopy volumes but also works with other high resolution imaging modalities. Detailed information of all original five datasets used in our evaluation is shown in Table 1.

**Table 1.**
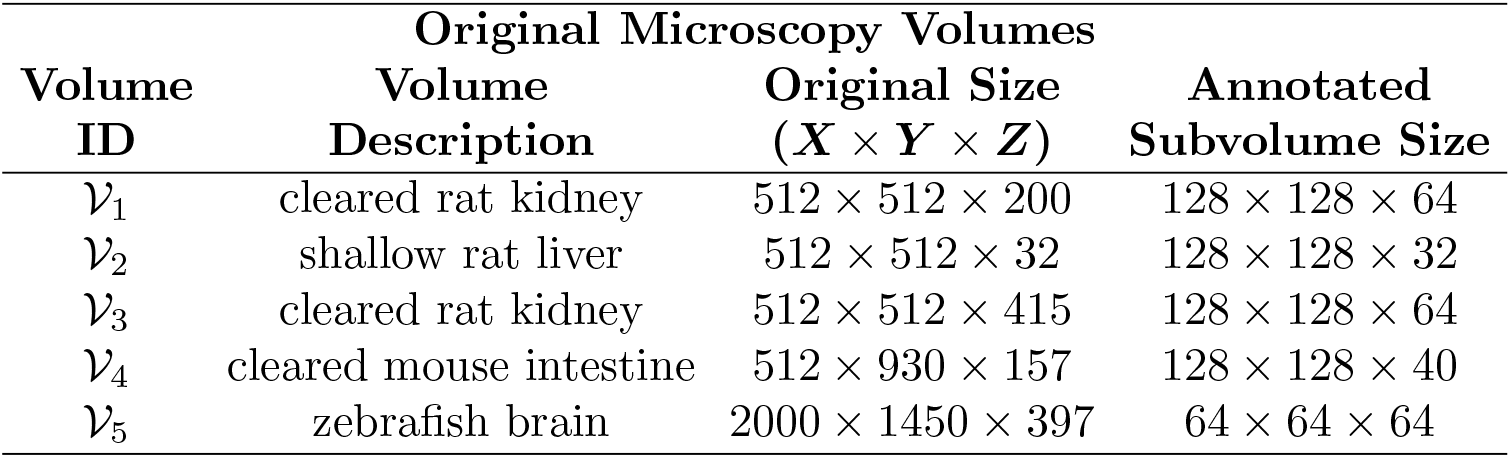
The description of the five datasets used for evaluation

| Volume ID | Original Microscopy Volumes |  |  |
| --- | --- | --- | --- |
| | Volume Description | Original Size ($X \times Y \times Z$) | Annotated Subvolume Size |
| $\mathcal{V}_1$ | cleared rat kidney | $512 \times 512 \times 200$ | $128 \times 128 \times 64$ |
| $\mathcal{V}_2$ | shallow rat liver | $512 \times 512 \times 32$ | $128 \times 128 \times 32$ |
| $\mathcal{V}_3$ | cleared rat kidney | $512 \times 512 \times 415$ | $128 \times 128 \times 64$ |
| $\mathcal{V}_4$ | cleared mouse intestine | $512 \times 930 \times 157$ | $128 \times 128 \times 40$ |
| $\mathcal{V}_5$ | zebrafish brain | $2000 \times 1450 \times 397$ | $64 \times 64 \times 64$ |

### 2.3 NISNet3D

Nuclei instance segmentation methods typically include a foreground and background separation step and a nuclei instance identification and segmentation step. Non-machine learning methods use traditional thresholding such as Otsu’s thresholding for separating background and foreground and use a watershed transformation to identify individual cell nuclei. However, classical thresholding methods are sensitive to variability in intensity and may not generate accurate segmentation masks. Also, watershed segmentation without accurate markers may generate under-segmentation or over-segmentation results. Machine learning methods typically generate more accurate results with enough training data. In addition, most current methods that are designed for nuclei instance segmentation only works on 2D images or uses a 2D to 3D reconstruction, which may not fully capture the 3D spatial information.

We improved upon these methods by designing a nuclei segmentation network, NISNet3D, using a modified 3D U-Net with attention and residual blocks, which can directly generate volumetric segmentation results. The modified 3D U-Net can be trained on both synthetic microscopy volumes and actual microscopy volumes if manual ground truth annotations are available. We generate more accurate markers than the direct distance transform used in classic watershed by learning a 3D vector field array which contains information that can be used to accurately infer the location of nuclei centroids as well nuclei boundaries. As shown in Figure 1, using the 3D vector field array, we further generate a 3D gradient array which contains gradient information, to aid in the detection of nuclei boundary and which also enables the generation and refinement of 3D markers that are to be used by a watershed method to perform segmentation. In addition, we provide two versions of NISNet3D: NISNet3D-slim and NISNet3D-syn that have the same network architectures but are trained with different datasets. NISNet3D-slim was trained only on a small number of synthetic/original volumes whereas NISNet3D-synth was trained using all the synthetic datasets we have. The details of NISNet3D-synth and NISNet3D-slim training and evaluation are described in Section 3.5. As demonstrated in Section 2.4 our methods achieve more accurate segmentation results.

### 2.4 Evaluation

We compared the NISNet3D with deep learning image segmentation methods including VNet [27], 3D U-Net [26], Cellpose [3], DeepSynth [6], and StarDist3D [15]. In addition, we also compare NISNet3D with several commonly used biomedical image processing tools including 3D Watershed [39], Squassh [40], CellProfiler [41], and VTEA [42]. We trained and evaluated the comparison methods using the same dataset as used for NISNet3D. We also used the same training and evaluation strategies as NISNet3D described in Section 3.5.

We use object-based metrics to evaluate nuclei instance segmentation accuracy. To reduce the bias [43], we use the mean Precision, mean Recall and mean F_1_ score on multiple IoU thresholds. We set *T*_IoUs_ = {0.25, 0.3, …, 0.45} for datasets 𝒱_1_-𝒱_4_, and set *T*_IoUs_ = {0.5, 0.55, …, 0.75} for datset 𝒱_5_. The selection of the *T*_IoUs_ is described in more detail below. We observed that the nuclei in datasets 𝒱_1_-𝒱_4_ are more challenging to segment than the nuclei in 𝒱_5_. If we use the same IoU thresholds for evaluating all the datasets, the evaluation accuracy for 𝒱_1_-𝒱_4_ will be much lower than the evaluation accuracy for 𝒱_5_ for all compared methods. Thus, we chose two different sets of IoU thresholds for 𝒱_1_-𝒱_4_ and 𝒱_5_, respectively.

We also examined commonly used object detection metrics: Average Precision (AP) [44, 45] by estimating the area under the Precision-Recall Curve [46] using the same thresholds for *T*_IoUs_ as described in the previous paragraph.

For example, AP_.25_ is the average precision with IoU threshold 0.25. In addition, we use the Aggregated Jaccard Index (AJI) [47] to integrate object and voxel errors.

All methods are optimized to achieve the best visual results by parameter tuning. This is further discussed in Section 3.5. The quantitative evaluation results for microscopy datasets 𝒱_1_-𝒱_4_ are shown in Table 2, and the quantitative evaluation results for microscopy dataset 𝒱_5_ are shown in Table 3 due to different IoU thresholds. We evaluate all subvolumes together for each dataset. In other words, for all subvolumes in a dataset, there is only one mP, mR, F_1_, AP, and mAP for each dataset. We noticed 3D Watershed, Squassh, Cellprofiler, and VTEA generate segmentation results with more heterogeneity for different subvolumes of each dataset. Figure 4 shows the AP scores using multiple IoU thresholds and the box plot of AJI on each subvolume of dataset _𝒱2_. The orthogonal views (XY focal planes and XZ focal planes) of the segmentation masks that are overlaid on the original microscopy subvolume for each method on _1_-_5_ are shown in Figure 5 and Figure 6. Note the colors correspond to different nuclei.

**Table 2.**
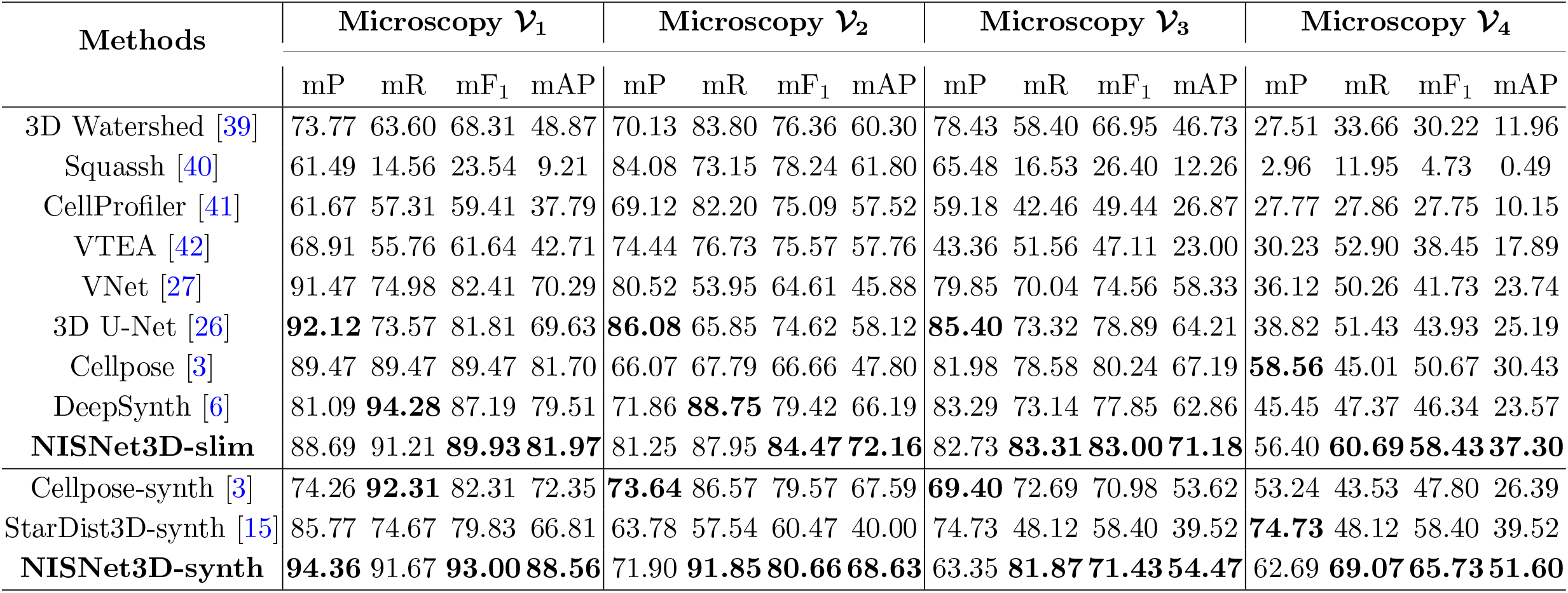
Comparison of object-based evaluation metrics for microscopy datasets 𝒱_1_ - 𝒱_4_. The best performance with respect to each metric are in bold

**Table 3.** Comparison of object-based evaluation results for microscopy dataset 𝒱_5_. The best performance with respect to each metric are in bold

| Methods | Microscopy $\mathcal{V}_5$ | | | | | | |
| --- | --- | --- | --- | --- | --- | --- | --- |
|  | mP | mR | mF<sub>1</sub> | AP<sub>.50</sub> | AP<sub>.75</sub> | mAP | AJI |
| StarDist3D [15] | 73.44 | 74.24 | 73.84 | 88.59 | 9.78 | 61.20 | 68.56 |
| DeepSynth [6] | 81.63 | 76.04 | 78.72 | 71.47 | 50.88 | 63.74 | 75.54 |
| Cellpose [3] | 96.15 | 94.47 | 95.30 | 94.90 | 83.82 | 91.21 | 81.31 |
| **NISNet3D-slim** | **96.89** | **96.24** | **96.56** | **95.98** | **88.84** | **93.62** | **83.90** |

**Fig. 4.**
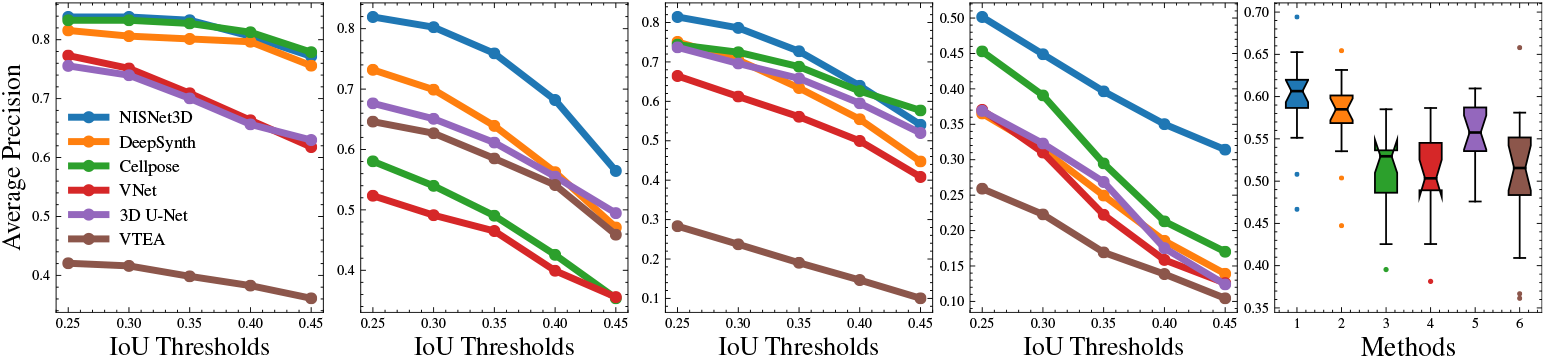
Evaluation results using Average Precision (AP) for multiple Intersection-over-Unions (IoUs) thresholds, *T*_IoUs_, for datasets 𝒱_1_-𝒱_4_, and box plots of Aggregated Jaccard Index (AJI) of each subvolume in dataset 𝒱_2_

**Fig. 5.**
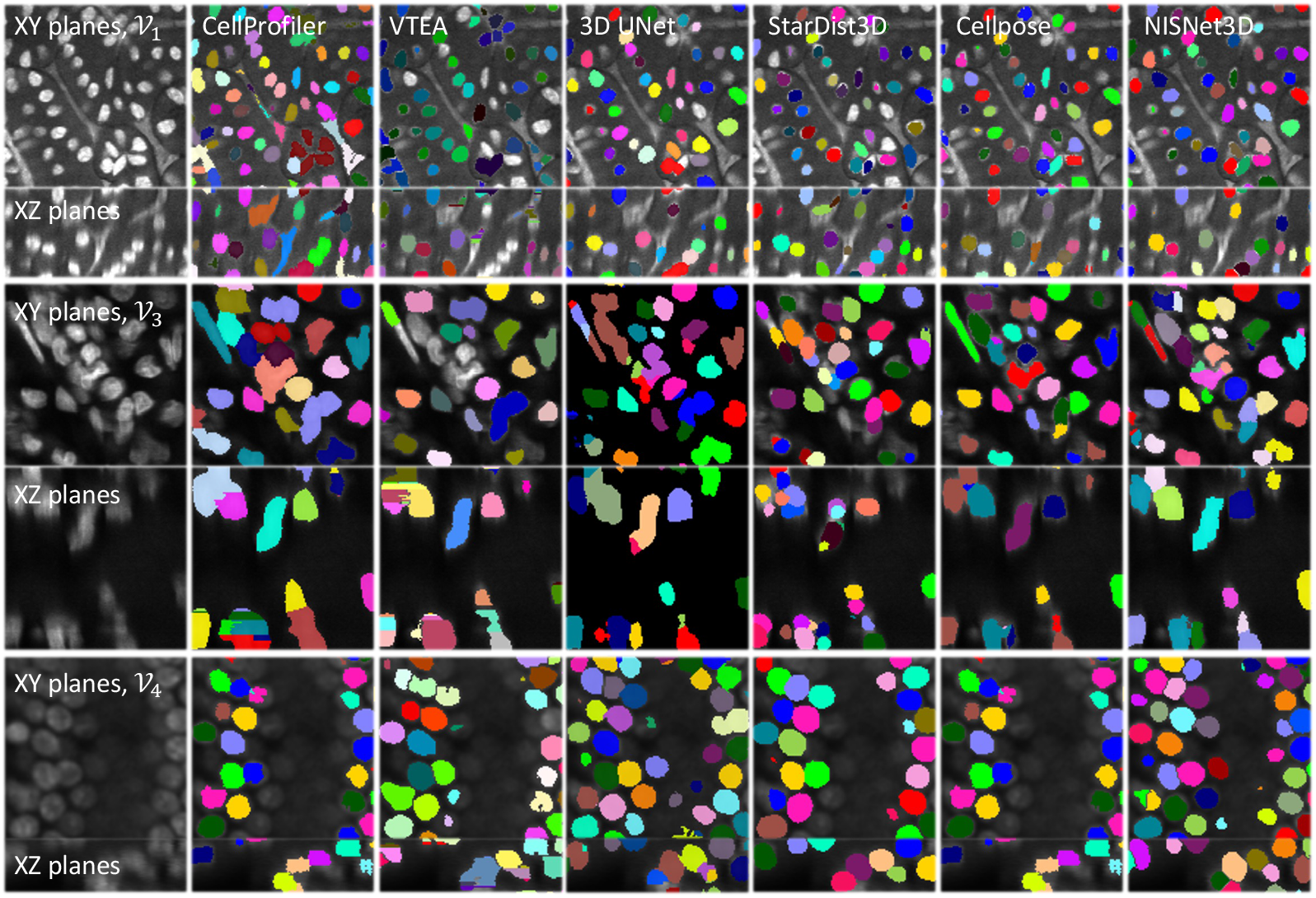
Visualization of segmentation results for XY and XZ focal planes

**Fig. 6.**
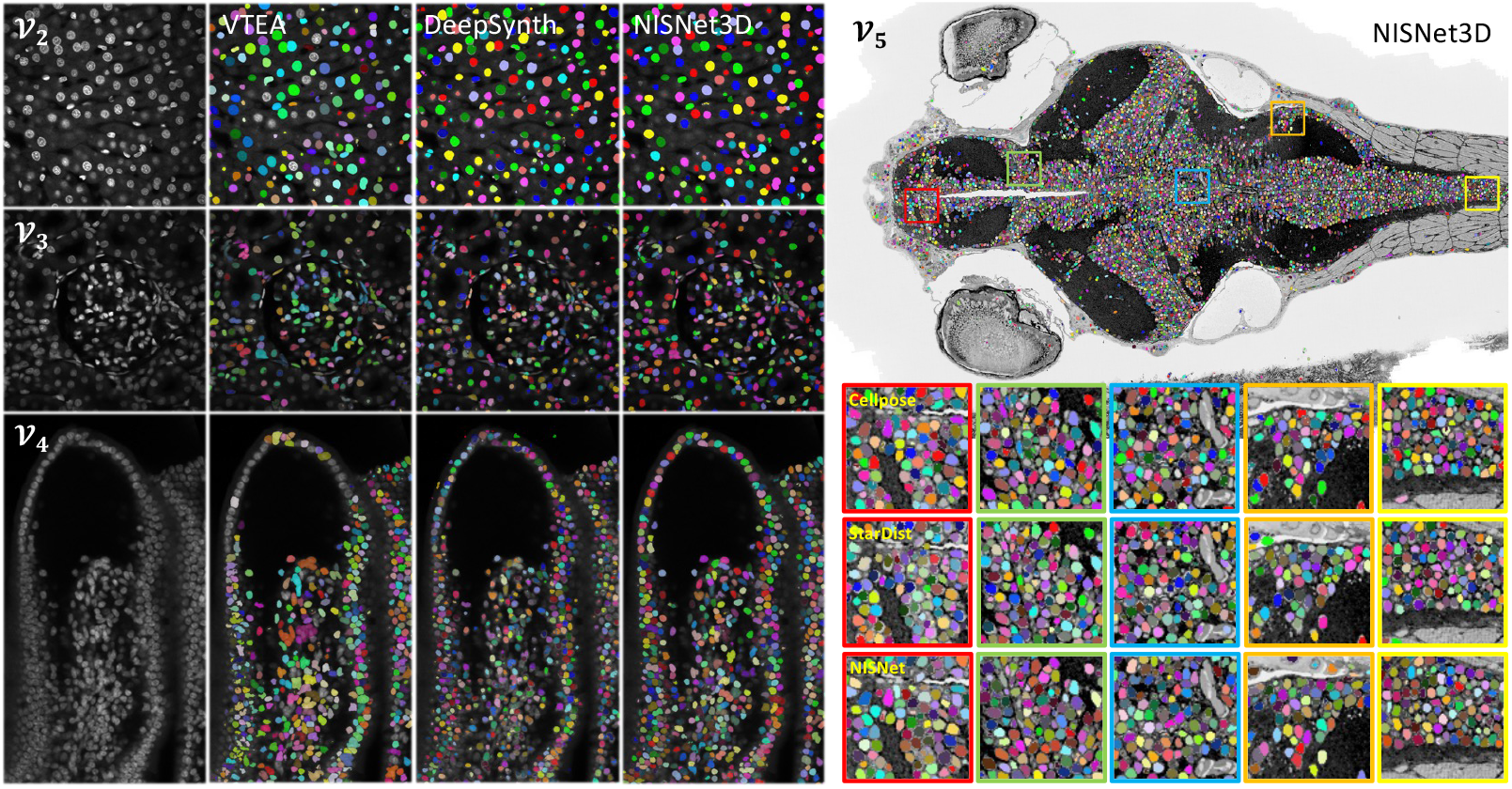
Visual comparison of the segmentations on a slice of the entire volume for datasets 𝒱_2_, 𝒱_3_, 𝒱_4_, and 𝒱_5_. Note that datasets 𝒱_2_, 𝒱_3_, and 𝒱_4_ are fluorescence microscopy volumes and dataset 𝒱_5_ is electron microscopy volume of zebrafish brain

### 2.5 Visualizing Differences

Entire microscopy volumes contain many regions with varying spatial characteristics. In order to see how the segmentation methods perform on various regions, Figure 7 shows the 2D to 3D reconstruction error from Cellpose and VTEA compared with NISNet3D. In addition, we use three methods for visualizing the differences between a “test segmented volume” and a “reference segmented volume”. The first method *Visualization Method A* shows the false negative voxels of a segmented volume, the second method *Visualization Method B* shows the undersegmentation regions where multiple nuclei in the “reference segmented volume” are detected as a single nucleus in the “test segmented volume”, and the second method *Visualization Method C* shows the false-negative objects by comparing with a “reference segmented volume”. The details of the three methods are described in Section 3.7. We use the NISNet3D segmented volume as the “reference segmented volume”. We visualize the segmentation differences between VTEA and NISNet3D and we also visualize the differences between DeepSynth and NISNet3D on entire volumes. Figure 8 shows the *Difference Volume* for VTEA, DeepSynth, and NISNet3D using *Visualization Method A* and *C*. Figure 9 shows the *Difference Volume* for using *Visualization Method B*.

**Fig. 7.**
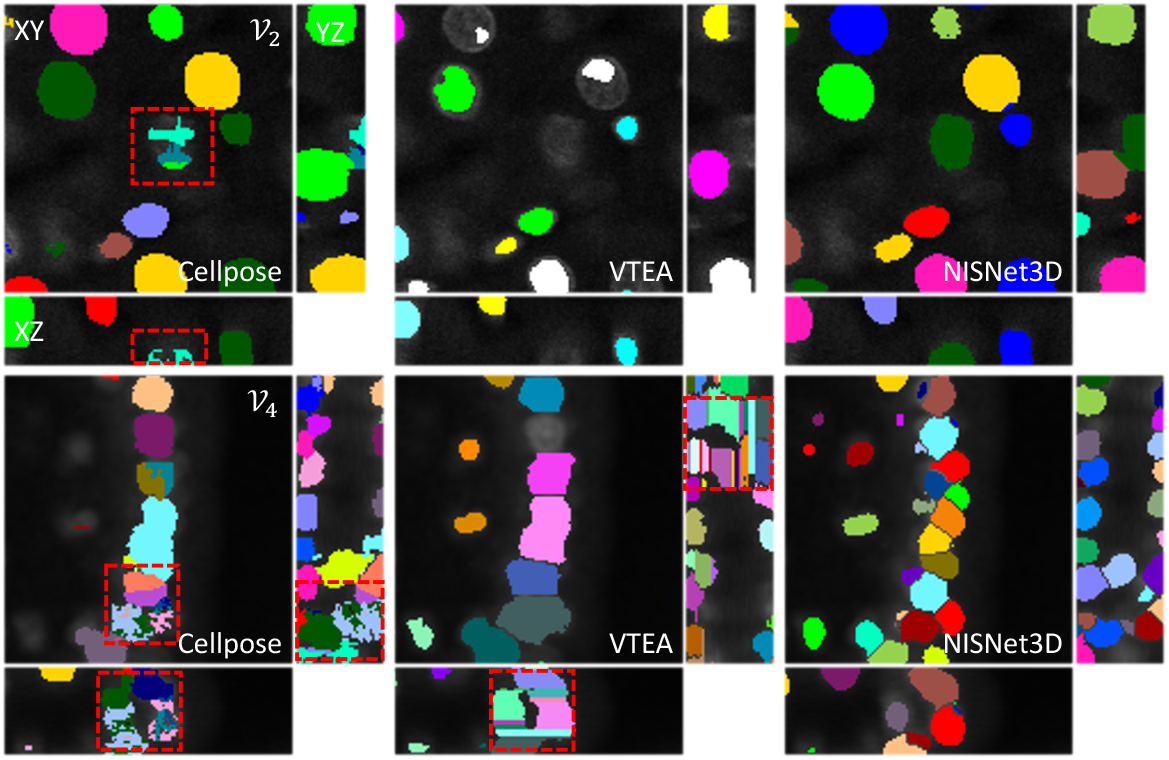
Visualization of segmentation results of Cellpose and VTEA compared with NIS-Net3D. The red boxes show the segmentation differences. Cellpose struggles reconstructing 3D segmentation from 2D segmentation results, leading to incomplete segmented 3D objects with holes when viewing from 3D perspective. VTEA struggles to distinguish different objects, resulting in under-segmentation. Also, the VTEA reconstruction results show strong artifacts for different slices of each 3D object.

**Fig. 8.**
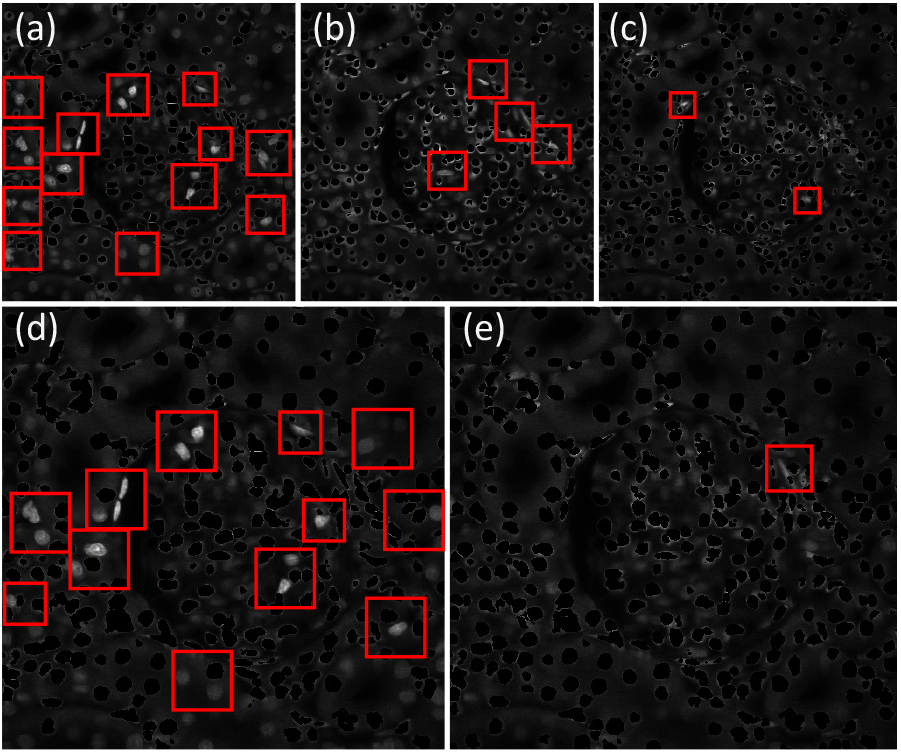
(a)(b)(c) shows the *Difference Volume* for VTEA, DeepSynth, and NISNet3D, respectively, using *Visualization Method A*. The nuclei voxels shown are the nuclei voxels in the original microscopy volume that are not segmented. (d)(e) shows the *Difference Volume* for VTEA and DeepSynth respectively, using *Visualization Method B*. The nuclei shown are the nuclei segmented by NISNet3D but completely missed by VTEA and DeepSynth, respectively. The red bounding boxes show the nuclei that are not segmented correctly

**Fig. 9.**
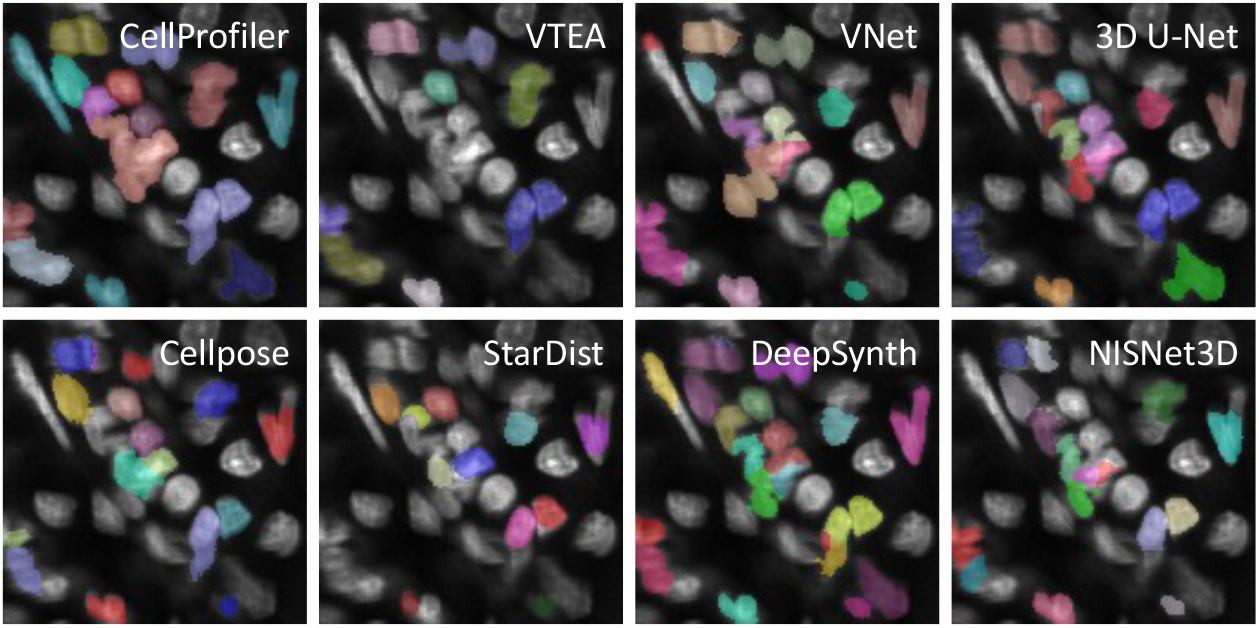
*Difference Volume* for dataset 𝒱_3_ using *Visualization Method B* which shows the undersegmentation regions where multiple nuclei in the “reference segmented volume” are detected as a single nucleus in the “test segmented volume”. The ground truth annotated volume is used as the “reference segmented volume” here

### 2.6 Discussion

In this paper, we describe a true 3D Nuclei Instance Segmentation Network, known as NISNet3D, for fluorescence microscopy images segmentation. Instead of segmenting 2D images slice by slice and reconstruct to 3D segmentation results, our approach directly works on 3D volumes by making use of a modified 3D U-Net and a nuclei instance segmentation system for separating touching nuclei. We improved the classical distance transform used for marker-controlled watershed by generating the 3D gradient volume and further obtain more accurate markers for each nuclei. Both visual and quantitative evaluation results demonstrate that directly using 3D CNN can improve the segmentation accuracy.

Due to the limited ground truth data, we use the synthetic microscopy volumes generated from SpCycleGAN for training. We improved the synthetic image generation by modeling nuclei as deformed ellipsoids. NISNet3D can be trained on both actual microscopy volumes and synthetic microscopy volumes generated using SpCycleGAN or a combination of both. NISNet3D can also be trained on synthetic data and further lightly retrained on limited number of other types of microscopy data as an incremental improvement. This can significantly reduce the computational resources needed for biologists since they only need an inference engine and update it with few annotated volumes. We demonstrate that NISNet3D performs well when compared to other methods on a variety of microscopy data both visually and quantitatively. To better inspect the segmentation results, we also present three visualization methods for visualizing segmentation differences in large 3D microscopy volumes without the need of ground truth annotations. Visualizing segmentation differences in different regions of a large microscopy volume is important for the understanding of model performance.

In the future, we will explore more synthetic nuclei segmentation masks generation methods that can better simulate the complex nuclei structures with more realistic density and distributions. We will extend the current SpCycleGAN to a 3D version such that it can directly generate 3D microscopy volumes with point spread function incorporated. In this way, the generated synthetic volumes will be more close to the distribution of real microscopy volumes, and can further improve the NISNet3D segmentation accuracy.

## 3 METHOD

The block diagram of proposed nuclei instance segmentation system is shown in Figure 10, which includes: (1) 3D microscopy image synthesis and annotated data, (2) NISNet3D training and inference, and (3) 3D nuclei instance segmentation.

**Fig. 10.**
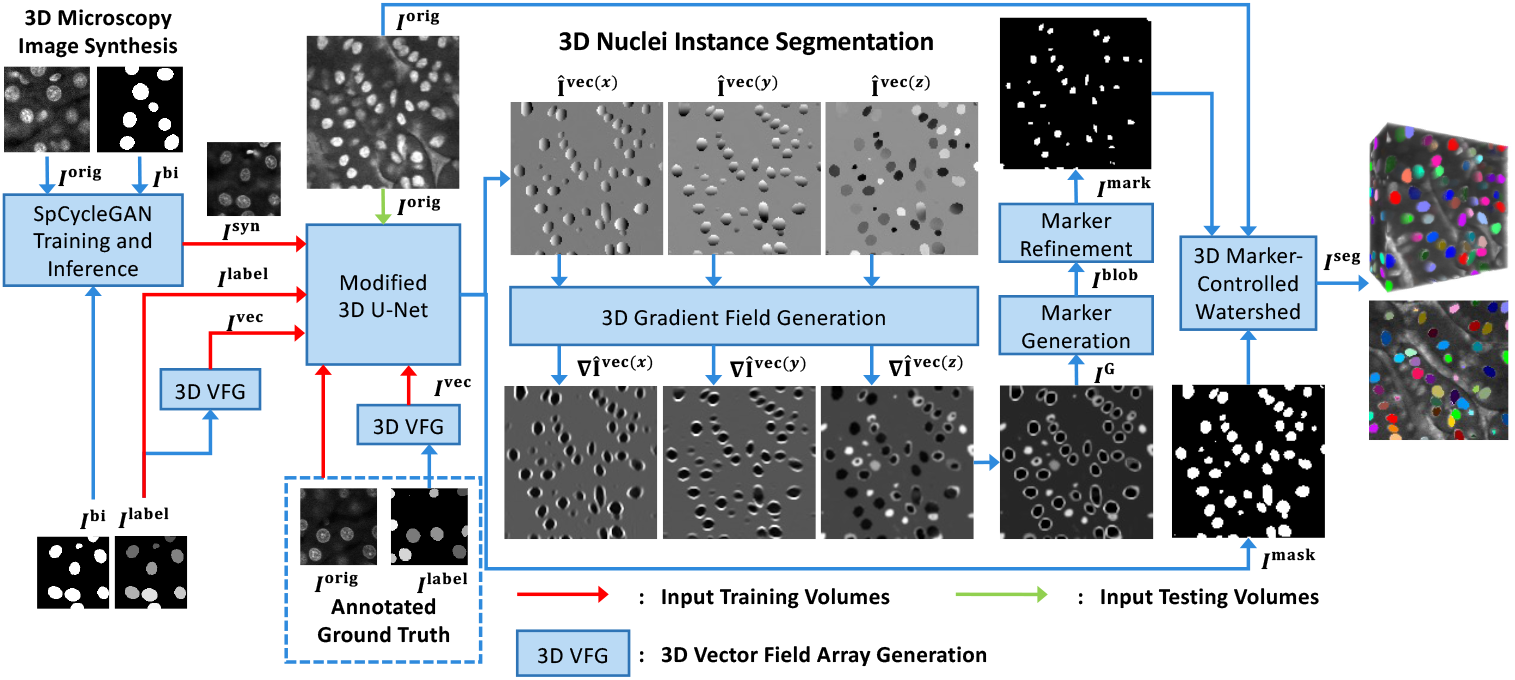
Overview of NISNet3D for 3D nuclei instance segmentation. Note: The training and synthetic image generation are also shown but are not explicitly part of NISNet3D. NISNet3D is trained with synthetic microscopy volumes generated from SpCycleGAN and/or with annotated actual volumes as indicated

### 3.1 Notation and Overview

In this paper, we denote a 3D image volume of size *X* ×*Y* ×*Z* voxels by *I*, and a voxel having coordinates (*x, y, z*) in the volume by *I*_(*x,y,z*)_. We will use superscripts to distinguish between the different types of volumes and arrays needed to achieve this goal. For example, *I*^orig^, *I*^bi^, and *I*^syn^ will be used to denote an original microscopy volume, a volume of binary segmentation masks, and a synthetic microscopy volume, respectively. The objective is to segment nuclei within *I*^orig^ using synthetic data. In addition, the various nuclei corresponding to the different masks in *I*^bi^ are marked with unique voxel intensities and comprise the volume *I*^label^. This is done to distinguish each nuclei instance and will be used during the training phase.

In addition to *I*^label^, NISNet3D generates from *I*^label^ a “vector field array,” *I*^vec^ of size *X* × *Y* × *Z* × 3, also for training purposes. Each element 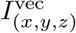 of *I*^vec^, located at (*x, y, z*), is a 3D vector 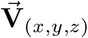 that is associated with the nucleus voxel 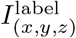 and points to the centroid of the nucleus with which it is associated. Detailed description of vector field generation will be discussed in Section 3.3.1.

For a given input volume, *I*^orig^, NISNet3D generates a corresponding volume *I*^mask^ of size *X* ×*Y* ×*Z* × 3 comprising the binary segmentation results as well as an estimate of the vector field array, *Î*_vec_, which is used to locate the nuclei centroids. In addition, *Î*_vec_ is used to generate a 3D array of gradients, *I*^G^ of size *X* ×*Y*×*Z*, that is used in conjunction with morphological operations and watershed segmentation to distinguish and separate touching nuclei as well as remove small objects. The final output, *I*^seg^, is a color-coded segmentation volume. Figure 10 provides an overview of our proposed approach.

### 3.2 3D Nuclei Image Synthesis

Deep learning methods generally require large amounts of training samples to achieve accurate results. However, manually annotating ground truth is a tedious task and impractical in many situations especially for 3D microscopy volumes. To address this, NISNet3D uses synthetic 3D microscopy volumes for training a segmentation network. It must be emphasized that NISNet3D can use synthetic volumes, annotated real volumes, or combinations of both. We demonstrate this in the experiments.

To generate synthetic volumes, we first generate synthetic segmentation masks that are used as ground truth masks, and then translate the synthetic segmentation masks into synthetic microscopy volumes using an unsupervised image-to-image translation model known as SpCycleGAN [7].

#### 3.2.1 Synthetic Segmentation Mask Generation

To generate a volume of synthetic binary nuclei segmentation masks, we iteratively add *N* initial binary nuclei to an empty 3D volume of size 128×128×128. Each initial nuclei is modeled as a 3D binary ellipsoid having random size, orientation, and location. The size, orientation, and location of the *n*^th^ ellipsoid are parameterized by ***a***^(*n*)^, ***θ***^(*n*)^, and ***t***^(*n*)^, respectively, where 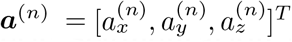 denotes a vector of semi-axes lengths of the *n*^th^ ellipsoid, 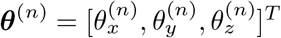 denotes a vector of rotation angles relative to the *x, y*, and *z* coordinate axes, and 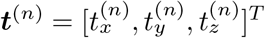 denotes the displacement/location vector of the *n*^th^ ellipsoid relative the origin. The parameters ***a***^(*n*)^, ***θ***^(*n*)^, and ***t***^(*n*)^ are randomly selected based on observations of nuclei characteristics in actual microscopy volumes.

Let 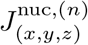 denote the *n*^th^ initial nucleus at location (*x, y, z*) having intensity/label *n* ∈ {1, …, N}, then

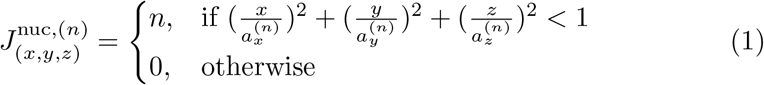

The ellipsoids are assigned different intensity values to differentiate them from each other. Subsequently, each 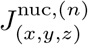 undergoes a random translation given by ***t***^(*n*)^ followed by random rotations specified in ***θ***^(*n*)^. Denoting the original coordinates by **X** and the translated and rotated coordinates by 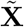, respectively then

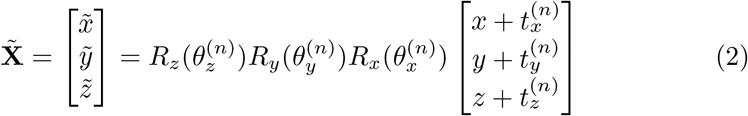

where 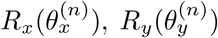 and 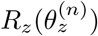 denote the rotation matrices relative to the *x, y, z* axes by angles 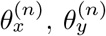, and 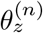, respectively. Thus, the final *n*^th^ ellipsoid 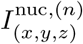 is given by 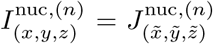 for all *n* ∈ {1, …, *N*}. We finally constrain the transformed nuclei such that they do not overlap by more than *t*_ov_ voxels. The ellipsoids 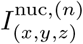, *n* ∈ {1, …, *N*}, are then used to fill in the binary volume *I*^bi^ of size *X*×*Y*×*Z*.

Since actual nuclei are not strictly ellipsoidal but look more like deformed ellipsoids (See Figure 2 (left column)), we use an elastic transformation [35] to deform *I*^bi^. To achieve this we first generate a “coarse displacement vector field”, *I*^coarse^, which is an array of size *d*×*d*×*d*×3, whose entries are independent and Gaussian distributed random variables having zero mean and variance *σ*^2^ (that is 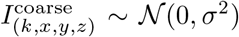)). The size parameter *d* is used to control the amount of deformation applied to the nuclei. *I*^coarse^ is then interpolated to size *X*×*Y*×*Z*×3, via spline interpolation [48] or bilinear interpolation [35] to produce a “smooth displacement vector field” *I*^smooth^.

The entry 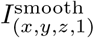 indicates the distance by which voxel 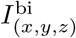 will be shifted and along the *x*-axis. Similarly, the elements 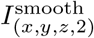 and 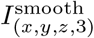 provide the distances by which 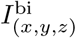 will be shifted along the *y* and *z*-axes, respectively.

In our experiments, we used spline interpolation and values for *d* ∈ {4, 5, 10}. Each element in *I*^coarse^ serves as an anchor point controlling the magnitude of elastic transform. Larger values of *d* result in larger deformations being applied through *I*^smooth^ whereas smaller *d* values produce less deformation. Examples of deformed ellipsoids are shown in Figure 2 (right column).

#### 3.2.2 Synthetic Microscopy Volume Generation

In the previous section we described how we generate binary segmentation masks. In the current section we describe how we use the masks to generate synthetic microscopy volumes. In particular, we use the unpaired image-to-image translation model known as SpCycleGAN [7, 36] for generating synthetic microscopy volumes. By unpaired we mean that the inputs to SpCycleGAN are volumes of binary segmentation masks and actual microscopy images that do not correspond with each other. In addition, the binary segmentation masks created above are not used as ground truth volumes for performing segmentation of actual microscopy images.

As shown in Figure 10, we use slices from volumes of the binary segmentation masks, *I*^bi,(*m*)^, *m* ∈ {1, …, *M*}, where *M* is the number of training samples/volumes, and slices from actual microscopy volumes *I*^orig,(*m*)^, *m* ∈{1, …, *M*} for training SpCycleGAN. After training we generate *L* synthetic microscopy volumes, *I*^syn,(*l*)^, *l* ∈ {1, …, *L* }, using synthetic microscopy segmentation masks, *I*^bi,(*l*)^, *l* ∈{1, …, *L*} other than the ones used for training. Note that since SpCycleGAN generates 2D slices, we use the slices to construct a 3D synthetic volume.

SpCycleGAN [7], an extension of CycleGAN [36], is shown in Figure 11. SpCycleGAN consists of two generators *G* and *F*, two discriminators *D*_1_ and *D*_2_. *G* learns the mapping from *I*^orig^ to *I*^bi^ whereas *F* performs the reverse mapping. Also, SpCycleGAN introduced a segmentor *S* for maintaining the spatial location between *I*^bi^ and *F*(*G*(*I*^bi^)). The entire loss function of SpCycleGAN is shown in Equation 3.

**Fig. 11.**
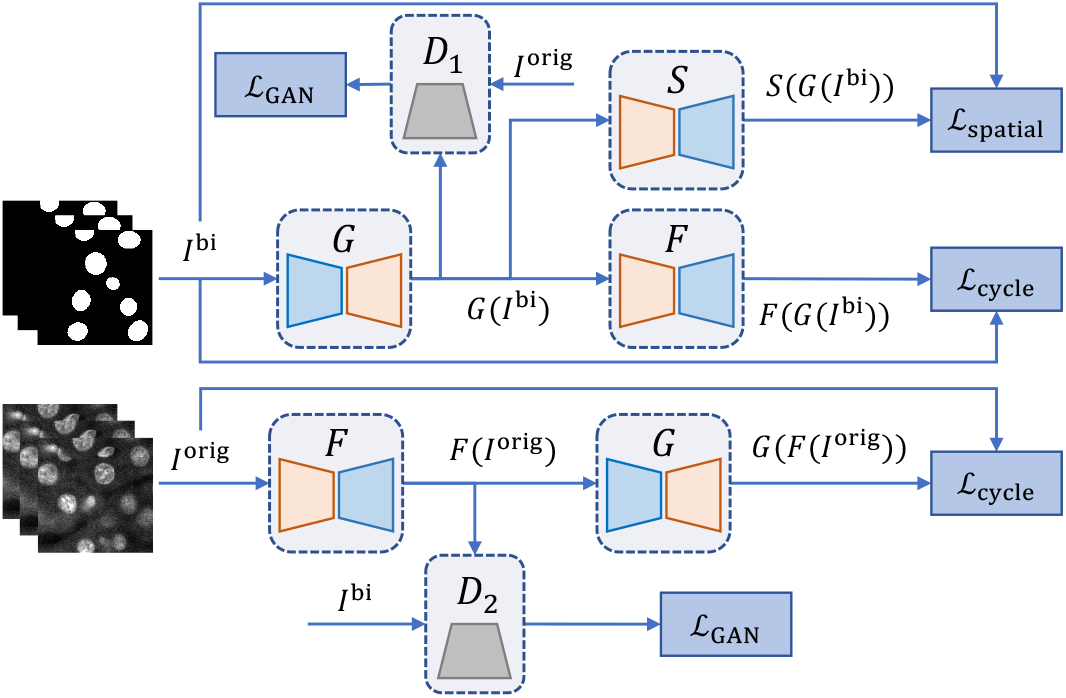
The architecture of SpCycleGAN that is used for generating synthetic microscopy volumes

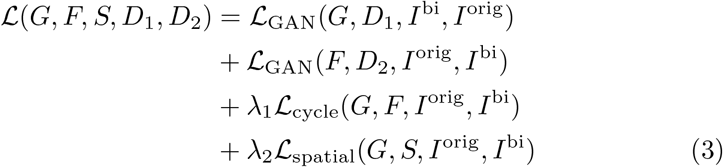

where *λ*_1_ and *λ*_2_ are weight coefficients controlling the loss balance between L_cycle_ and L_spatial_, and

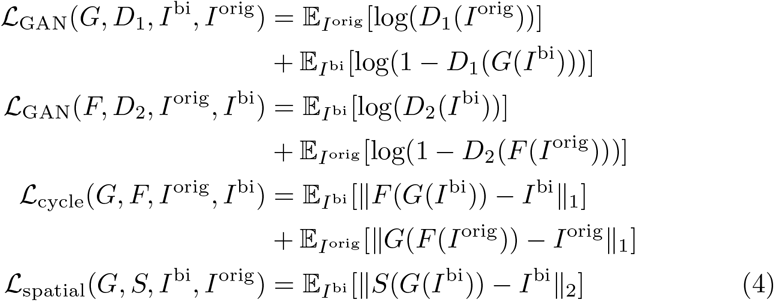

where ∥·∥_1_ and ∥·∥_2_ denotes the *L*_1_ norm and *L*_2_ norm, respectively. E_*I*_ is the expected value over all the input volumes of a batch to the network.

### 3.3 NISNet3D

NISNet3D is based on a modified 3D U-Net architecture. In this section, we describe the architecture, how to train and inference using the modified 3D U-Net, as well as nuclei instance segmentation performed by NISNet3D. Figure 12 provides an overview of NISNet3D.

**Fig. 12.**
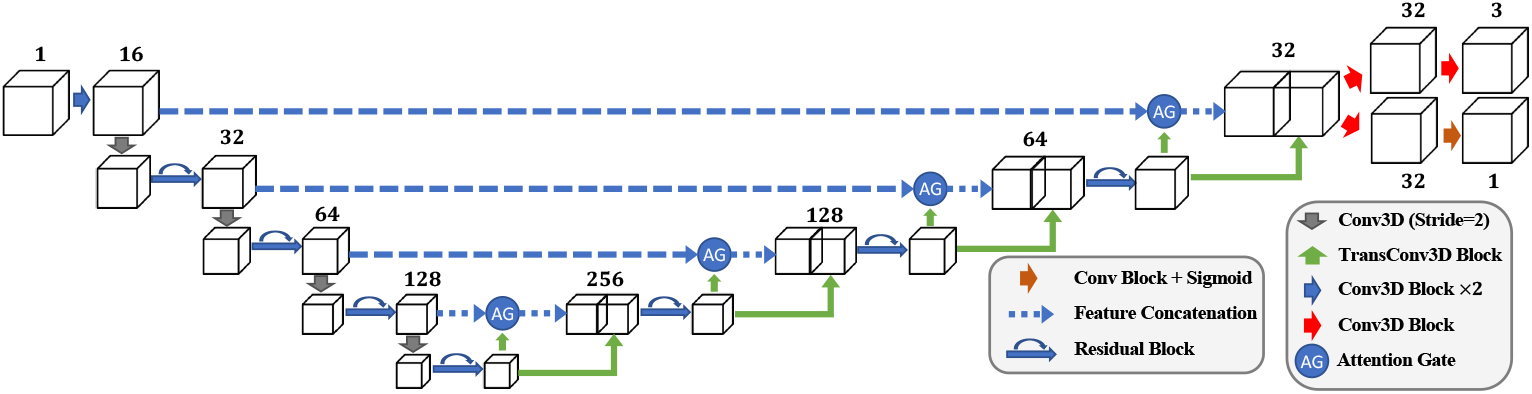
The proposed NISNet3D uses a modified 3D U-Net architecture with residual blocks, attention gates, and multi-task learning module

#### 3.3.1 Modified 3D U-Net

As mentioned previously NISNet3D utilizes a modified 3D U-Net. In this section we describe its architecture given in Figure 12. In general NISNet3D employs an encoder-decoder network (See Figure 12) which outputs the same size array as the input. The encoder consists of multiple Conv3D Blocks (Figure 13(a)) and Residual Blocks (Figure 13(b)) [49]. Instead of using max pooling layers, we use Conv3D Blocks with stride 2 for feature down-sampling, which introduces more learnable parameters. Each convolution block consists of a 3D convolution layer with filter size 3 × 3 × 3, a 3D batch normalization layer, and a leaky ReLU layer. The decoder consists of multiple TransConv3D blocks (Figure 13(d)) and attention gates (Figure 13(c)). Each TransConv3D block includes a 3D transpose convolution with filter size 3 × 3 × 3 followed by 3D batch normalization and leaky ReLU. We use a self-attention mechanism described in [50] to refine the feature concatenation while reconstructing the spatial information.

**Fig. 13.**
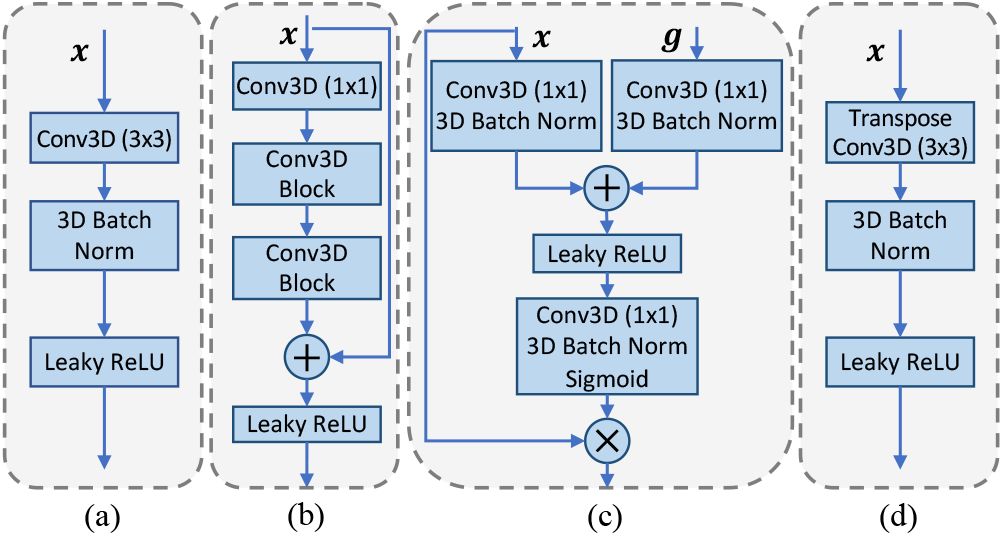
(a) Conv3D Block, (b) Residual Block, (c) Attention Gate, (d) TransConv3D Block

##### Training The Modified 3D U-Net

The modified 3D U-Net can be trained on both synthetic microscopy volumes *I*^syn^ or actual microscopy volumes *I*^orig^ if manual ground truth annotations are available.

As shown in Figure 10, during training, NISNet3D’s modified 3D U-Net takes an *I*^syn^ or *I*^orig^, an *I*^label^, and an *I*^vec^ as input. *I*^label^ is the *X*×*Y*×*Z* grayscale label volume corresponding to *I*^bi^ where different nuclei are marked with unique pixel intensities. It used is used by NISNet3D to learn the segmentation masks. Moreover, to facilitate segmentation NISNet3D identifies nuclei centroids and estimates their locations using a “vector field,” *I*^vec^. In particular, *I*^vec^ is part of the ground truth data used during training to enable the network to learn nuclei centroid locations. In addition, the 3D vector field array helps identify the boundaries of adjacent/touching nuclei since their corresponding vectors tend to point in different directions.

As indicated previously a “vector field,” *I*^vec^, is an *X* × *Y* × *Z* × 3 array whose elements are 3D vectors that point to the centroids of the corresponding nuclei. Specifically, 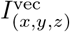 is a 3D vector at location (*x, y, z*) and points to the nearest nucleus centroid. We also denote the *X* × *Y* × *Z* array consisting of the first component of each 3D vector by *I*^vec(*x*)^. Similarly, we denote the arrays of the second and third components by *I*^vec(*y*)^ and *I*^vec(*z*)^, respectively, that is 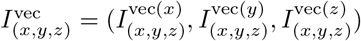.

The first step in the 3D vector field array generation (VFG) (see in Figure 14) is to obtain the centroid of each nucleus in *I*^label^. We denote the *k*^th^ nucleus as the set of voxels with intensity *k* in *I*^label^, and the location of the centroid of the *k*^th^ nucleus by (*x, y, z*). We define 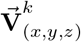 to be the 3D vector from (*x, y, z*) to (*x*_*k*_, *y*_*k*_, *z*_*k*_) and set 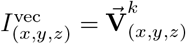 as long as the voxel at (*x, y, z*) is not a background voxel. Note that if a voxel 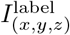 is a background voxel (i.e. 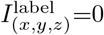), the corresponding entry 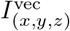 in the vector field array is set to **0**, as shown in Equation 5.

**Fig. 14.**
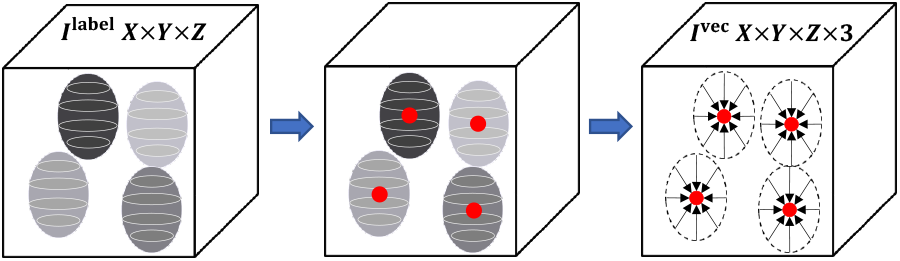
Steps for 3D vector field array generation (VFG). Each nucleus voxel in the 3D vector field array, *I*^vec^ represents a 3D vector that points to the centroid of current nucleus

Using *I*^syn^ or *I*^orig^, *I*^label^, and *I*^vec^, NISNet3D then outputs an estimate of the 3D vector field array *Î*_vec_ (*Î*_vec(*x*)_, *Î*_vec(*y*)_, *Î*_vec(*z*)_) and the 3D binary segmentation mask volume *I*^mask^.

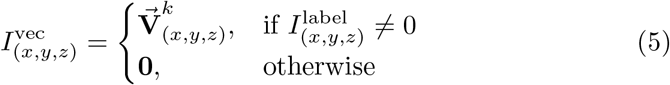

##### Loss Functions

Given *I*^syn^ or *I*^orig^, *I*^label^, and *I*^vec^, the modified 3D U-Net simultaneously learns the nuclei segmentation masks *I*^mask^ and the 3D vector field array *Î*_vec_. In the modified 3D U-Net two branches but no sigmoid function are used to obtain *Î*_vec_ because the vector represented at a voxel can point to anywhere in an array. In other words, the entries in *Î*_vec_ can be negative numbers or large numbers. Unlike previous methods [31, 51, 52] that directly learn the distance transform map, the 3D vector field array contains both the distance and direction of the nearest nuclei centroid from the current voxel location. This can help NISNet3D avoid the multiple detection of irregular shaped nuclei. The output 3D vector field array *Î*_vec_ is compared with the ground truth vector field array *I*^vec^ and the error between them minimized using the Mean Square Error (MSE) loss function. Similarly, the segmentation result *I*^mask^ is compared with the ground truth binary volume *I*^bi^ and the difference minimized using a combination of the Focal Loss [53] ℒ _FL_ and Tversky Loss [54] ℒ _TL_ metrics. Denoting a ground truth binary volume *I*^bi^ by *S*, the corresponding segmentation result *I*^mask^ by *ŝ*, a ground truth vector field array *I*^vec^ by *V*, and the estimated vector field array *Î*_vec_ by 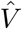, then, the entire loss function is,

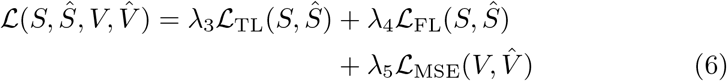

where

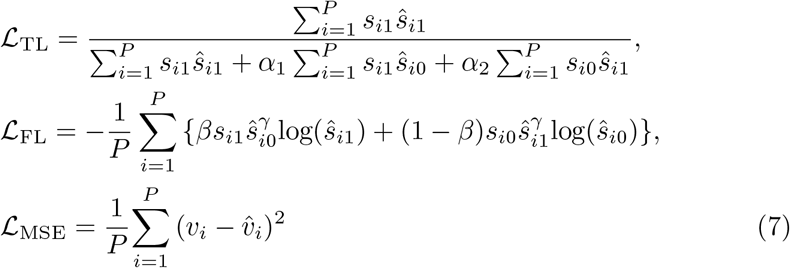

where α_1_ + *α*_2_ = 1 are two hyper-parameters in Tversky loss [54] that control the balance between false positive and false negative detections. *β* and *γ* are two hyper-parameters in Focal loss [53] where *β* balances the importance of positive/negative voxels, and *γ* adjusts the weights for easily classified voxels. *v*_*i*_ ∈ *V* is the *i*^th^ element of *V*, and 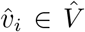 is the *i*^th^ entry in 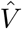. Similarly, *s*_*i*_ ∈ *S* is the *i*^th^ voxel in *S*, and *ŝ*_*i*_ ∈ *ŝ* is the *i*^th^ voxel in *ŝ*. We define *ŝ*_*i*0_ to be the probability that the *i*^th^ voxel in *ŝ* is a nuclei, and *ŝ*_*i*1_ as the probability that the *i*^th^ voxel in *ŝ* is a background voxel. Similarly, *s*_*i*1_ = 1 if *s*_*i*_ is a nuclei voxel and 0 if *s*_*i*_ is a background voxel, and vice versa for *s*_*i*0_. Lastly, *P* is the total number of voxels in a volume.

##### Modified 3D U-Net Inference

To segment a large microscopy volume, we propose an divide-and-conquer inference scheme shown in Figure 15. We use an inference window of size *K* × *K* × *K* that slides along each original microscopy volume *I*^orig^ of size *X* × *Y* × *Z* and crops a subvolume. Considering that cropping may result in some nuclei being partially included, and that these partially included nuclei which lie on the border of the inference window may cause inaccurate segmentation results, we construct a padded window by symmetrically padding each cropped subvolume by 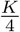 voxels on each border.

**Fig. 15.**
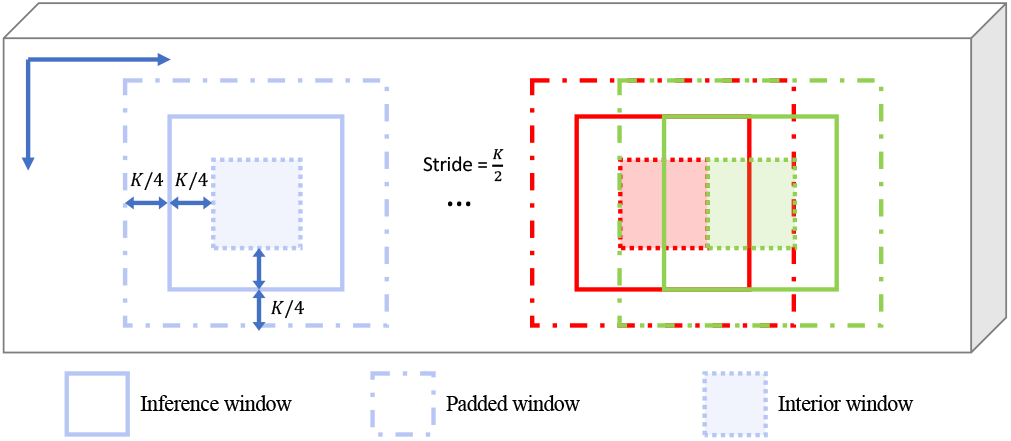
Proposed divide-and-conquer inference scheme for segmenting large microscopy volumes

Also, the stride of the moving window is set to 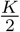 so with every step it slides, it will have 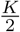 voxels overlapping with the previous window. For the inference results of every *K* ×*K* ×*K* window, only the interior and centered 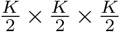 subvolume, which we denoted as the interior window, will be used as the segmentation results. In this paper, we use *K* = 128 for all testing data. Once the inference window slides along the entire volume, a segmentation volume *I*^mask^ and 3D vector field array *Î*_vec_ of size *X* × *Y* × *Z* will be generated. In this way, we can inference on any size input volume, especially very large volumes.

#### 3.3.2 3D Nuclei Instance Segmentation

Based on the output of the modified 3D U-Net, a 3D gradient field and an array of markers are generated and used to separate densely clustered nuclei. Moreover, the markers are refined to attain better separation. As mentioned previously, each element/vector in the estimated 3D vector field array *Î*_vec_ point to the centroid of the nearest nucleus. Most often, the vectors associated with voxels on the boundary of two touching nuclei point in different directions that correspond to the locations of the centroids of the touching nuclei, and hence have a large difference/gradient. We employ such larger gradients to detect the boundaries of touching and also overlapping nuclei.

Let Δ*Î*_vec_ denote the gradient of *Î*_vec_ which is obtained as given in Equation 8:

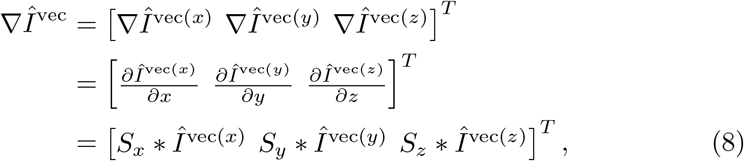

where *Î*_vec(*x*)_, *Î*_vec(*y*)_, and *Î*_vec(*z*)_ are the *x, y*, and *z* sub-arrays of the estimated 3D vector field array *Î*_vec_ respectively, *S*_*x*_, *S*_*y*_, *S*_*z*_ are 3D Sobel filters, and * is the convolution operator. We also define the gradient map *I*^G^ as the maximum of the gradient components along the *x, y* and *z* directions (see Equation 9).

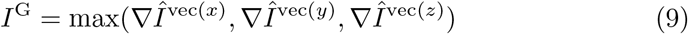

Elements of *I*^G^ that correspond to boundaries of touching nuclei have larger values (larger gradients) which can be used to identify individual nuclei. We subsequently threshold *I*^G^ via a thresholding function *τ*(*x, T*_m_) where *τ*(*x, T*_m_) = 1 if *x*≥*T*_m_ otherwise 0. This step is used to highlight and distinguish boundary voxles of individual nuclei from spurious edges that may arise from finding the gradients. The value of *T*_m_ can affect the number of voxels delineated as boundary voxels. Larger values of *T*_m_ result in thinner boundaries whereas smaller values of *T*_m_ result in a thicker nuclei boundaries. We then subtract the result from the binary segmentation mask *I*^mask^ obtained from the modified 3D U-Net, and then apply that to a non-linear half wave rectifier/RELU activation function *σ*(·) that sets all negative values to 0. We denote the result as *I*^blob^ as given in Equation 10. The difference *I*^mask^ −*τ*(*I*^G^, *T*_m_) removes boundary voxles of touching nuclei while emphasizing the interior. Since the difference can be negative in some cases, we use *σ*(·) to eliminate negative values. Hence, *I*^blob^ highlights the interior regions of nuclei which maybe used as markers to perform watershed segmentation [39, 55].

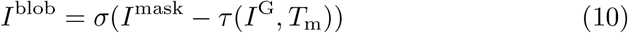

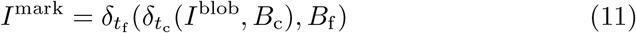

However, in some cases *I*^blob^ may still contain boundary information. To refine it we use 3D conditional erosion with a coarse structuring element *B*_c_ and a fine structuring *B*_f_ as shown in Figure 16. To achieve this we first identify the connected components [56] in *I*^blob^ and then iteratively erode each connected component using *B*_c_ until its size is smaller than some threshold *t*_c_. We then continue eroding each connected component using *B*_f_ until its size is smaller than another threshold *t*_f_. We denote the final result by *I*^mark^, as shown in Equation 11, where 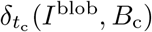 defines iterative erosion of all connected component in *I*^blob^ by a coarse structuring element *B*_c_ until the size of each component is smaller than the coarse object threshold *t*_c_. Similarly, 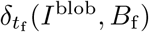, denotes iterative erosion by a fine structuring element, *B*_f_, until the size of the object is smaller than the fine object threshold *t*_f_. Finally, marker-controlled watershed [39] is used to generate instance segmentation masks *I*^seg^ using *I*^mark^. Small objects in *I*^seg^ that are less than 20 voxels in size are then removed, and each object color coded for visualization.

**Fig. 16.**
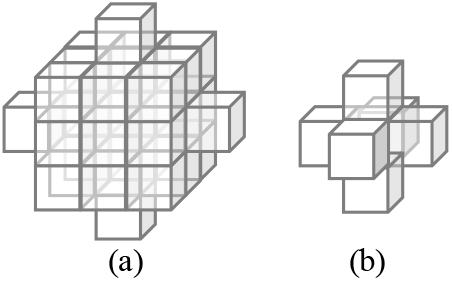
(a) coarse 3D structuring element *B*_c_ and (b) fine 3D structuring element *B*_f_ for 3D conditional erosion

### 3.4 Comparison Methods

We compared NISNet3D with several deep learning image segmentation methods including VNet [27], 3D U-Net [26], Cellpose [3], DeepSynth [6], and StarDist3D [15]. In addition, we also compare NISNet3D with several commonly used biomedical image analysis tools including 3D Watershed [39], Squassh [40], CellProfiler [41], and VTEA [42].

VNet [27] and 3D U-Net [26] are two popular 3D encoder-decoder networks with shortcut concatenations designed for biomedical image segmentation. Cellpose [3] uses a modified 2D U-Net for estimating image segmentation and spatial flows, and uses a dynamic system [3] to cluster pixels and further separate touching nuclei. When segmenting 3D volumes, Cellpose works from three different directions slice by slice and combines the 2D segmentation results into a 3D segmentation volume [3]. DeepSynth uses a modified 3D U-Net to segment 3D microscopy volumes and uses watershed to separate touching nuclei [6]. Similarly, StarDist3D uses a modified 3D U-Net to estimate the star-convex polyhedra used to represent nuclei [15].

Alternative non deep-learning based techniques such as 3D Watershed [39] uses the watershed transformation [55] and conditional erosion [39] for to perform nuclei segmentation. Similarly, Squassh, an ImageJ plugin for both 2D and 3D microscopy image segmentation, uses active contours [40]. CellProfiler is an image processing toolbox and provides customized image processing and analysis modules [41], whereas VTEA is an ImageJ plugin that combines various approaches including Otsu’s thresholding and watershed to segment 2D nuclei slice by slice and reconstruct the results into a 3D volume [42].

For comparison purposes, we trained and evaluated the methods above using the same dataset as used for NISNet3D. We also used the same training and evaluation strategies used for NISNet3D as described in Section 3.5. We provide a discussion of the results in Section 2.6. Note that 3D Watershed, Squassh, CellProfiler, and VTEA do not need to be trained because they use more traditional image analysis techniques.

### 3.5 Experimental Settings

The parameters used for generating *I*^bi^ are provided in Table 4 where (*a*_min_, *a*_max_) is the range of the ellipsoid semi-axes lengths, *t*_ov_ is the maximum allowed overlapping voxels between two nuclei, and *N* is the total number of nuclei in a synthetic volume. These parameters are based on visual inspection of nuclei characteristics in actual microscopy volumes.

**Table 4.** The training evaluation strategies for NISNet3D and comparison methods

| Parameters For Generating Synthetic Microscopy Volumes |  |  |  |  |  |  | NISNet3D-slim Training Strategies |  | NISNet3D-slim Evaluation |
| --- | --- | --- | --- | --- | --- | --- | --- | --- | --- |
| Volume Name | $a_{\min}$ | $a_{\max}$ | $t_o$ | $N$ | $d$ | $\sigma$ | Version | Training | |
| $\mathcal{V}_1$ | 4 | 8 | 5 | 500 | 10 | 1 | $\mathcal{M}_1$ | 50 volumes of synthetic $\mathcal{V}_1$ | Tested on 1 subvolume of $\mathcal{V}_1$ |
| $\mathcal{V}_2$ | 10 | 14 | 10 | 70 | 4 | 1 | $\mathcal{M}_2$ | 50 volumes of synthetic $\mathcal{V}_2$ | Tested on 16 subvolumes of $\mathcal{V}_2$ |
| $\mathcal{V}_3$ | 6 | 10 | 200 | 200 | 5 | 2 | $\mathcal{M}_3$ | 50 volumes of synthetic $\mathcal{V}_3$ | 3-fold cross-validated on 9 subvolumes of $\mathcal{V}_3$ |
| $\mathcal{V}_4$ | 8 | 10 | 100 | 560 | 4 | 4 | $\mathcal{M}_4$ | lightly retrained using $\mathcal{M}_3$ on 1 subvolume of $\mathcal{V}_4$ | Tested on 4 subvolumes of $\mathcal{V}_4$ |
| $\mathcal{V}_5$ | - | - | - | - | - | - | $\mathcal{M}_5$ | 27 subvolumes of $\mathcal{V}_5$ | Tested on 27 subvolumes of $\mathcal{V}_5$ |

SpCycleGAN was trained on unpaired *I*^bi^ and *I*^orig^ and the trained model was used to generate synthetic microscopy volumes. The weight coefficients of the loss functions were set to *λ*_1_ = *λ*_2_ = 10 based on the experiments described in [7]. The synthetic microscopy volumes were verified by a biologist (one of the co-authors).

For 3D Watershed, we used a 3D Gaussian filter to preprocess the images and used Otsu’s method to segment/isolate objects from the background of each volume. Subsequently the 3D conditional erosion described in Section 3.3.2 was used to obtain the markers, and the marker-controlled 3D watershed method, which was implemented by the Python Scikit-image library, was used to separate touching nuclei.

In the case of CellProfiler [41], customized image processing modules including inhomogeneity correction, median filtering, and morphological erosion were used to preprocess the images, and the default “IdentifyPrimaryObject” module was used to obtain 2D segmentation masks for each slice. The 2D segmentations were then merged into a 3D segmentation volume using the blob-slice method described in [42, 57].

With regards to Squassh [40], we used “Background subtraction” with a manually tuned “rolling ball window size” parameter. The rest of the parameters were set to their default values. We observed that Squassh did fairly well on volume 𝒱_2_ but totally failed on 𝒱_4_ due to the densely clustered nuclei. For VTEA [42], a Gaussian filter and background subtraction were used to preprocess the image. The object building method was set to “Connect 3D”, and the segmentation threshold determined automatically. We manually tuned the parameters “Centroid offset”, “Min vol”, and “Max vol” to obtain the best visual segmentation results. Finally, watershed was utilized to perform segmentation. Since VTEA and Cellprofier’s “IdentifyPrimaryObject” modules only work on 2D images, we see that their segmentation results, shown in Figure 5, suffer from over-segmentation errors in the XZ planes.

As far as VNet [27], 3D U-Net [26] and DeepSynth [6] are concerned, we improved the segmentation results by using our 3D conditional erosion described in Section 3.3.2 followed by 3D marker-controlled watershed to separate touching nuclei. In the case of Cellpose, we used the “nuclei” style and since the training of Cellpose is only limited to 2D images, we trained Cellpose on every XY focal planes of our subvolumes following the training schemes given in Table 4. We observe that Cellpose has trouble capturing some very large or small nuclei in an input subvolume and performs worse on “thinner” subvolumes containing more non-ellipsoidal nuclei. Figure 7 shows the 2D to 3D reconstruction errors for Cellpose and VTEA when compared with32 NISNet3D’s results. For StarDist, we observed that it has difficulty segmenting concave objects or objects that are not ellipsoidal or star like in shape (non-star-convex objects) in 𝒱_3_ and achieves better performance on regular ellipsoidal nuclei in 𝒱_4_ (See Figure 5).

Both SpCycleGAN and NISNet3D were implemented using PyTorch. We used 9-block ResNet for the generators *G, F*, and the segmentor *S* (see Figure 11). The discriminators *D*_1_ and *D*_2_ (Figure 11) were implemented with the “PatchGAN” classifier [58]. SpCycleGAN was also trained with an Adam optimizer [59] for 200 epochs with an initial learning rate 0.0002 that linearly decays to 0 after the first 100 epochs. Figure 3 depicts the generated synthetic nuclei segmentation masks and corresponding synthetic microscopy images.

As mentioned above, we provide two versions of NISNet3D: NISNet3D-slim and NISNet3D-syn for different application scenarios. We trained 5 versions of NISNet3D-slim, denoted as ℳ_1_-ℳ_5_, using three training methods (See Table 4). First, we trained three models ℳ_1_, ℳ_2_, and ℳ_3_ using the synthetic versions of the three volumes 𝒱_1_-𝒱_3_, respectively. Next, we transfer the weights from ℳ_3_ and continue training on a limited number of actual microscopy subvolumes of 𝒱_4_ to produce the fourth model M_4_. Finally, we directly train ℳ_5_ on subvolumes of actual microscopy data 𝒱_5_ only. After training, we evaluate the models ℳ_1_-ℳ_5_ through two different evaluation schemes. In one method we directly test the models ℳ_1_, ℳ_2_, ℳ_4_, and ℳ_5_ using all the subvolumes of the actual microscopy volumes 𝒱_1_, 𝒱_2, 4_, and 𝒱_5_, respectively, since they were not used for training. In the case of ℳ_3_, we use 3-fold cross-validation to split the actual microscopy volume 𝒱_3_ into 9 subvolumes that are randomly divided into 3 equal sets and then iteratively update the model on one of the sets and test on other two sets. We used cross-validation to show the effectiveness of our method when the evaluation data is limited. Note that when lightly retraining ℳ_3_ we update all its parameters while continue to train on actual microscopy volumes. The training and evaluation scheme for all models is provided in Table 4.

NISNet3D-synth, on the other hand is trained on 800 synthetic microscopy volumes that include the synthetic versions of 𝒱_1_-𝒱_4_, and tested separately on each manually annotated original microscopy datasets described in Table 1. NISNet3D-synth is designed for the situation where no ground truth annotations are available. In contrast, NISNet3D-slim is used for the case where limited ground truth annotated volumes are available. In this case synthetic volumes are used for training and a small amount of actual ground truth data is for used for light retraining.

Both NISNet3D-slim and NISNet3D-synth were trained for 100 epochs using the Adam optimizer [59] with a constant learning rate of 0.001. The weight coefficients of the loss function were set to *λ*_3_ = 1, *λ*_4_ = 10, *λ*_5_ = 10, the hyper-parameters *β, γ* of ℒ_FL_ were set to 0.8 and 2, respectively, and the hyper-parameters *α*_1_, *α*_2_ in ℒ_TL_ were set to 0.3 and 0.7 [54, 60], respectively. The nuclei instance segmentation parameters used in our experiments are provided in Table 5.

**Table 5.**
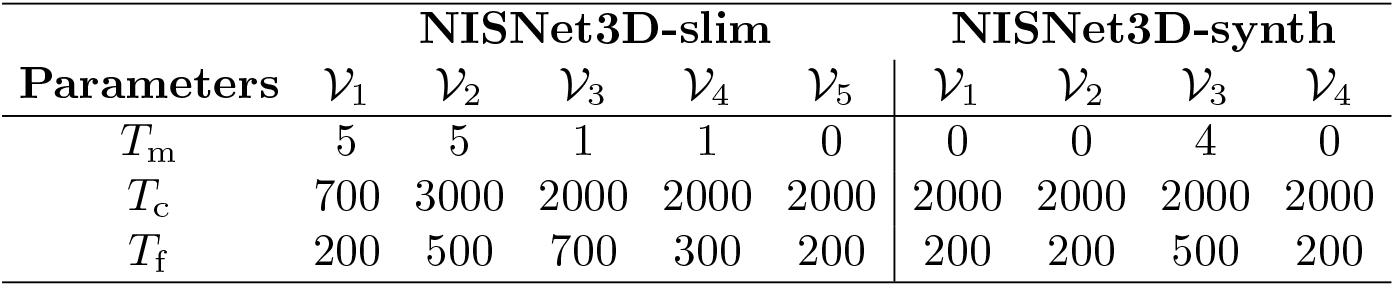
Parameters used for NISNet3D nuclei instance segmentation

### 3.6 Evaluation Metrics

We use object-based metrics to evaluate nuclei instance segmentation accuracy. To this end, we define 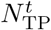 as the number of True Positive detections when the Intersection-over-Union (IoU) [61] between a detected nucleus and a ground truth nucleus is greater than some threshold of *t* voxels. Similarly, 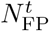 is the number of False Positives greater than *t* voxels, and 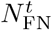 is the number of False Negatives [62, 63] greater than *t* voxels, respectively. 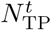 measures how many nuclei in a volume are correctly detected, and the higher the value of 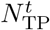 the more accurate the detection. 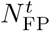 represents the detected nuclei that are not actually nuclei but are false detections, and 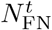 represents the number of nuclei that were not detected. A precise and accurate detection method should have high 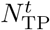 but low 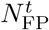 and 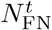 values.

Based on 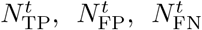 we define the following metrics that are used to reduce bias [43]: mean Precision 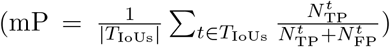 mean Recall 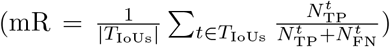, and mean F_1_ score 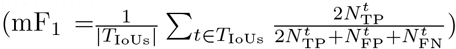 that are the mean of Precision, Recall, and the F_1_ score, respectively, over a set of multiple IoU thresholds *T*_IoUs_. We set *T*_IoUs_ = {0.25, 0.3, …, 0.45} for datasets 𝒱 _1_-𝒱 _4_, and set *T*_IoUs_ = {0.5, 0.55, …, 0.75} for datset 𝒱_5_. In addition, we obtained the Average Precision (AP) [44, 45] (a commonly used object detection metric), by estimating the area under the Precision-Recall Curve [46], using the same sets of thresholds in *T*_IoUs_. For example, AP_.25_ is the average precision evaluated at an IoU threshold of 0.25. The mean Average Precision (mAP) is then obtained as 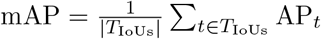.

The reason for using different thresholds *T*_IoUs_ for 𝒱 _1_-𝒱 _4_ compared to those used for 𝒱 _5_ is that throughout the experiments we observed that the nuclei in datasets 𝒱 _1_-𝒱 _4_ are more challenging to segment than the nuclei in _5_. Using the same IoU thresholds for evaluating all the datasets resulted in a lower evaluation accuracy for the volumes 𝒱 _1_-𝒱 _4_ than for V_5_. Thus, we chose two different sets of IoU thresholds for 𝒱 _1_-𝒱 _4_, and 𝒱 _5_, respectively.

Finally, we use the Aggregated Jaccard Index (AJI) [47] to integrate object and voxel errors. The AJI is defined as:

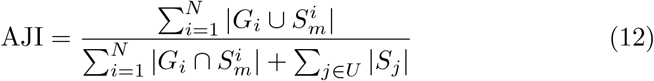

where *G*_*i*_ denotes the *i*th nucleus in the ground truth volume *G* having a total number of *N* nuclei, *U* is the volume of segmented nuclei without corresponding ground truth, and 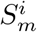 is the *m*th connected component in the segmentation mask which has the largest Jaccard Index/similarity with *G*_*i*_. Note that each segmented nucleus with index *m*, that is belongs *m*th connected component, cannot be used more than once.

The values of all the parameters for each method were chosen to achieve the best visual results, as discussed in Section 2.6. The values of the various metrics used are given in Table 2 and Table 3 for each microscopy dataset. Figure 4 shows the AP scores using multiple IoU thresholds and the box plot of AJI for each subvolume of the dataset 𝒱 _2_. The orthogonal views (XY focal planes and XZ focal planes) of the segmentation masks are overlaid on the original microscopy subvolume for each method on 𝒱 _1_-𝒱 _5_ as shown in Figure 5 and Figure 6. Note that the different colors correspond to different nuclei.

### 3.7 Visualizing the Differences

Entire microscopy volumes contain many regions with varying spatial characteristics. In order to see how the segmentation methods perform on various regions, we propose three methods for visualizing the errors and the differences between a “test segmented volume” and a “reference segmented volume”. We use the NISNet3D segmented volume as the reference segmented volume to visualize the segmentation differences between VTEA and NISNet3D, and also between DeepSynth and NISNet3D for the entire volumes. The three methods employed will be referred to as *Visualization Method A, B*, and *C*, respectively, and use an *Overlay Volume* and three types of *Difference Volumes*. Note that *Visualization Method A* does not need a “reference segmented volume” whereas *Visualization Method B* and *C* need a “reference segmented volume”. Next we describe how to generate an *Overlay Volume*. Using the notation shown in Figure 10, we denote an original microscopy volume as *I*^orig^ and *m* as the maximum intensity of *I*^orig^. For a “reference segmented volume” (obtained via NISNet3D), we denote the binary segmentation masks for *I*^orig^ as *I*^mask^ and denote the color-coded segmentation for *I*^orig^ as *I*^seg^. The R, G, and B components of *I*^seg^ are denoted by 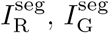, and 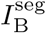, respectively. Similarly, for a “test segmented volume” (obtained via any other method), we denote *C*^bi^ as the binary segmentation masks for *I*^orig^, and denote *C* as the colorcoded segmentation of *I*^orig^ with R, G, and B components *C*_R_, *C*_G_, and *C*_B_, respectively.

We denote the *Overlay Volume* as *L* having R, G, and B components *L*_R_, *L*_G_, and *L*_B_, respectively. The *Overlay Volume* for a “test segmented volume” is generated by adding the original microscopy volume to each of the R, G, and B components of the color-coded segmented volume, as described in Equation 13.

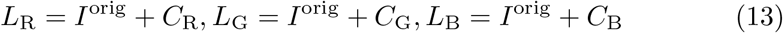

Let 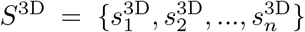 be the set of all 3D segmented nuclei in *C*, where 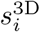 is a volume having the same size as *C* but only contains the *i*th segmented 3D nucleus, and let 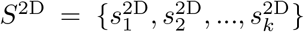 be the set of all 2D objects in *C* from each XY focal plane, where 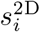 is a volume of same size as *C* but only contains the *i*th segmented 2D nucleus. Similarly, let 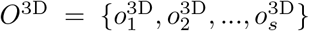 be the set of all 3D objects in *I*^seg^, and let 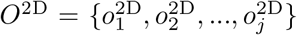 be the set of all 2D objects in *I*^seg^ from each slice. We use *S*^3D^ and *O*^3D^ in our generation of the three types of *Difference Volumes* using the *Visualization Methods A, B*, and *C*.

#### 3.7.1 Visualization Method A

The *Difference Volume* generated by *Visualization Method A* shows voxels in the original microscopy volume that are not segmented by either test or reference methods. The input to *Visualization Method A* is the original microscopy volume and a segmented volume (“test segmented volume” or “reference segmented volume”). Using VTEA as an example: a VTEA segmented volume is weighted by the maximum intensity *m*, subtracted from the original microscopy volume, and negative differences set to 0 as indicated in Equation 14.

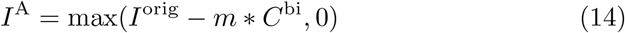

This operation produces *Difference Volume, I*^A^, which highlights the voxels in the original microscopy image that are not segmented. We replace *C*^bi^ in Equation 14 with *I*^mask^ to obtain the *Difference Volume* for the “reference segmented volume” and NISNet3D. Figure 8 ((a), (b), (c)) shows the *Difference Volumes* generated by *Method A* for VTEA, DeepSynth, and NISNet3D using dataset 𝒱_3_.

#### 3.7.2 Visualization Method B

The *Difference Volume, I*^B^, generated by *Visualization Method B* emphasizes the under-segmentation regions, that is the regions where multiple nuclei in the “reference segmented volume” are detected as a single nucleus in the “test segmented volume”. Here we use NISNet3D as “reference segmented volume” and use VTEA or DeepSynth as “test segmented volume”. The input to *Visualization Method B* is a VTEA (or DeepSynth) segmented volume and a NISNet3D segmented volume.

Using VTEA as an example: if two or more nuclei in the NISNet3D segmented volume intersect with the same single nucleus in the VTEA segmented volume, we emphasize the single nucleus segmented by VTEA in *I*^B^, using Equation 15.

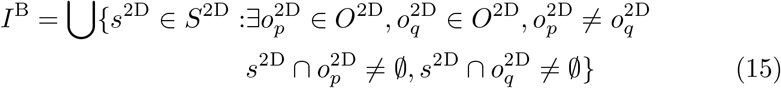

The resulting volume *I*^B^ is overlaid on the original microscopy volume using Equation 13. Figure 9 depicts the *Difference Volumes* generated by *Visualization Method B* on a subvolume of dataset V_3_.

#### 3.7.3 Visualization Method C

The *Difference Volume, I*^C^, generated by *Visualization Method C* highlights nuclei segmented by a “reference segmented volume” but are completely missed by a “test segmented volume”. The input to *Visualization Method C* is a VTEA (or DeepSynth) segmented volume and a NISNet3D segmented volume. Using VTEA as an example: if the voxels of a nucleus in NISNet3D segmented volume do not intersect with any voxels of any segmented nucleus from the VTEA segmented volume, then *I*^C^ will emphasize this nucleus from NISNet3D, using Equation 16:

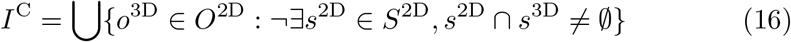

The *Difference Volumes* for *Visualization Method A* and *C* are shown in Figure 8.

## 4 ACKNOWLEDGMENT

This research was partially supported by a George M. O’Brien Award from the National Institutes of Health under grant NIH/NIDDK P30 DK079312 and the endowment of the Charles William Harrison Distinguished Professorship at Purdue University.

The authors have no conflicts of interest.

The original image volumes used in this work were provided by Malgorzata Kamocka, Sherry Clendenon, and Michael Ferkowicz at Indiana University. We gratefully acknowledge their cooperation.

The NISNet3D source code package is available upon request to. The source code is released under Creative Commons License Attribution-NonCommercial-ShareAlike - CC BY-NC-SA. The source code cannot be used for commercial purposes. The test volumes are also available at [37].

## Notes

### Competing Interest Statement

The authors have declared no competing interest.

### Summary of Updates

The introduction has been rewritten. The entire paper has been reorganized to better describe the methods.

